# CD4-mediated immunity shapes neutrophil-driven tuberculous pathology

**DOI:** 10.1101/2024.04.12.589315

**Authors:** Benjamin H Gern, Josepha M Klas, Kimberly A Foster, Sara B Cohen, Courtney R Plumlee, Fergal J Duffy, Maxwell L Neal, Mehnaz Halima, Andrew T Gustin, Alan H Diercks, Alan Aderem, Michael Gale, John D Aitchison, Michael Y Gerner, Kevin B Urdahl

## Abstract

Pulmonary *Mycobacterium tuberculosis* (Mtb) infection results in highly heterogeneous lesions ranging from granulomas with central necrosis to those primarily comprised of alveolitis. While alveolitis has been associated with prior immunity in human post-mortem studies, the drivers of these distinct pathologic outcomes are poorly understood. Here, we show that these divergent lesion structures can be modeled in C3HeB/FeJ mice and are regulated by prior immunity. Using quantitative imaging, scRNAseq, and flow cytometry, we demonstrate that Mtb infection in the absence of prior immunity elicits dysregulated neutrophil recruitment and necrotic granulomas. In contrast, prior immunity induces rapid recruitment and activation of T cells, local macrophage activation, and diminished late neutrophil responses. Depletion studies at distinct infection stages demonstrated that neutrophils are required for early necrosis initiation and necrosis propagation at chronic stages, whereas early CD4 T cell responses prevent neutrophil feedforward circuits and necrosis. Together, these studies reveal fundamental determinants of tuberculosis lesion structure and pathogenesis, which have important implications for new strategies to prevent or treat tuberculosis.

## Introduction

The outcomes of aerosol infection with *Mycobacterium tuberculosis* (Mtb), the bacteria that causes tuberculosis (TB), are highly heterogeneous and shaped by prior immunity, including immunity from vaccination or prior Mtb exposure.^1^ The pulmonary granuloma, an organized aggregate of immune cells, often with a necrotic core that destroys normal lung architecture, is frequently considered the hallmark lesion of TB.^2^ However, human post-mortem studies in the pre-antibiotic era showed that many Mtb-infected lung lesions do not exhibit a granulomatous architecture.^3^ In primary TB, where patients had no previous exposure to Mtb, pulmonary lesions usually start as granulomas, with a core of macrophages that often undergo necrosis centrally surrounded by a lymphocytic cuff.

Conversely in post-primary TB (when individuals had prior Mtb exposure), lesions usually first developed into pneumonia-like alveolitis, with infected macrophages contained within intact alveolar sacs infiltrated by lymphocytes.^1^ Despite appreciation of the association between prior immunity and Mtb lesion types for more than a century, the mechanisms by which prior immunity promotes the development of alveolitis instead of granulomas remain unknown.

In modern times, human post-mortem studies are rare and most research dissecting TB immunity is performed in animals without prior Mtb exposure. These studies have revealed many insights about the varied microenvironments within the granuloma that restrict immune function, including distinct myeloid cell niches (macrophage subtypes, monocytes, granulocytes) and various factors that suppress T cell effector functions. T cells are frequently relegated to the peripheral cuff, being unable to infiltrate the granuloma cores and engage in cognate interactions with infected cells.^4,5^ Lesions also directly suppress T cells through local immunoregulatory factors, including TGFβ^6^ and products of tryptophan metabolism,^4^ some of which are spatially partitioned in distinct immunoregulatory domains leading to localized immune suppression.^7^ It is largely unknown how these microenvironments and immune regulatory factors differ in granulomatous versus alveolitis lesions, raising the possibility that host-directed therapies may be effective only in certain lesion types. Furthermore, necrotic lesions can progress to lung-destructive cavitary TB disease, which takes longer to respond to antibiotic treatment and has a high risk of relapse and recurrent infection. Thus, understanding how to prevent these types of lesions from forming could lead to new strategies to curb severe manifestations of disease.^8–12^

Historically, mouse models have lacked the ability to dissect relationships between lesion structure and disease control. Mtb-infected C57BL/6 mice, the most commonly used mouse strain for TB research due to the abundance of tools for mechanistic studies, do not form necrotic granulomas when infected with a conventional aerosol dose of 50-100 colony-forming units (CFU).^13^ However, recent work has shown that reducing the infectious dose to a more-physiologic 1-3 CFU results in well-circumscribed lesions that share properties with stereotypical human lesions, including discrete and segregated regions containing T cells, infected macrophages, and B cell follicles, respectively.^14^ Furthermore, C3HeB/FeJ mice do develop necrotizing granulomas, especially when infected with hypervirulent Mtb strains of the W-Beijing lineage.^15^ A single gene that confers the extreme susceptibility and necrotic lesions of C3HeB/FeJ mice has been identified as *Sp140*, an epigenetic regulator with chromatin-binding domains that can influence inflammatory gene transcription,^16^ and *Sp140*^-/-^ mice on a C57BL/6 background exhibit similar TB susceptibility and pathology phenotypes as C3HeB/FeJ mice.^17^ Use of these mouse models has led to the identification of critical signaling pathways that regulate Mtb infection outcomes, including type I IFN vs IL-1, as well as insights into the temporal processes driving inflammation and disease: neutrophil recruitment, cellular death, pDC sensing, and IFN production/signaling.^17,18^ In addition, studies in collaborative-cross mice have shown that host genetics can heavily influence disease susceptibility and ability to control Mtb after immunization with bacillus Calmette– Guérin (BCG).^19–21^ Protection in these models was associated with differences in T cell effector responses and concordant structural changes of pulmonary lesions, suggesting that ability of immune cells to deliver their critical effector functions within the lesions may be associated with improved outcome.

Here, we report a mouse model that recapitulates these two divergent types of TB lesions, necrotizing granulomas in non-immune animals and alveolitis in those with prior immunity. Using advanced immunologic techniques and quantitative spatial approaches, we find that pre-existing immunity results in enhanced early T cell and macrophage activation at infected sites, which is associated with decreased neutrophil clustering and tissue destruction at late timepoints. Using depletion studies, we further show that CD4 T cells are critical for the protection afforded by pre-existing immunity against necrosis, and in their absence all lesions develop necrosis and increased neutrophil infiltration. Conversely, we find that neutrophils are required for lesion necrosis throughout infection, including both the early generation and propagation of centralized necrosis, and result in reduced T cell and macrophage activation. Together, these studies provide insight into protective immunity afforded by pre-existing immunity and reveal pivotal opposing roles for CD4 T cells and neutrophils in driving disease outcomes.

## Results

## Pre-existing immunity abrogates the formation of necrotic granulomas

To examine the impact of pre-existing or ongoing immune responses against Mtb on de novo lesion structure development and disease progression, we utilized the C3HeB/FeJ (C3H) mouse model, which generates large necrotic granulomas akin to those found in human primary TB, especially after aerosol infection with hypervirulent or high transmission W-Beijing Mtb strains.^15,22^ To induce pre-existing or concomitant immunity to Mtb, we used two established modalities: subcutaneous BCG immunization 8 weeks prior to Mtb challenge, as well as concomitant Mtb infection (CoMtb), in which a low-level chronic Mtb infection is established in the cutaneous lymph node after intradermal Mtb inoculation.^23,24^ Mice administered either BCG or CoMtb, or unimmunized controls, were aerosol infected with a conventional dose (CD, 50-100 CFU) of SA161 Mtb, a hypervirulent clinical isolate from the W-Beijing lineage. As expected, when assessed at day 98 (d98) post-infection (p.i.), unimmunized mice developed large granulomas with a central necrotic core rimmed by foamy macrophages, surrounded by a lymphocytic cuff, as well as multiple smaller lesions without overt necrosis (Fig 1A). In stark contrast, both CoMtb and BCG completely blocked the formation of necrotic granulomas, instead inducing smaller, less-organized lesions comprised of histiocytes and lymphoid cells, and exhibiting less neutrophil infiltration (Fig 1A, S1A). Blinded, quantitative evaluation of multiple lung pathology metrics using principal component analysis (PCA) showed that CoMtb resulted in the greatest overall changes in pathology as compared to Mtb infected control mice (Fig 1B, S1B, Table S1). Both BCG and CoMtb markedly reduced bacterial burdens at d28 p.i., together indicating that pre-existing and concomitant immunity offer robust protection at early timepoints. Differences in lung CFU were less pronounced 98 days post-infection, although the bacterial burdens were still significantly lower in the CoMtb group (Fig 1C). Given the improved protection seen with CoMtb as compared to BCG, we chose CoMtb as the modality of prior immunity to dissect mechanistically.

**Figure 1:**
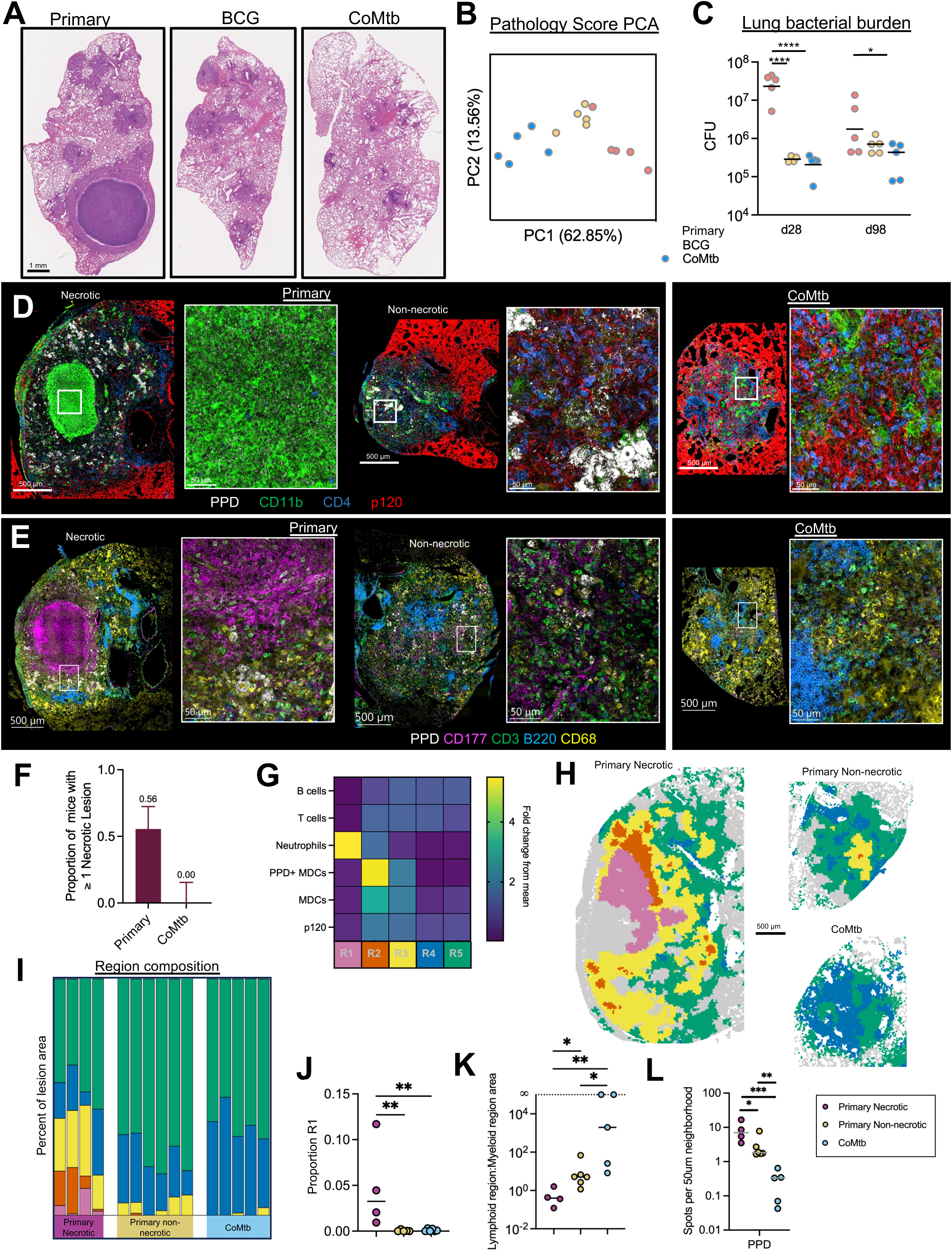
Pre-existing immunity abrogates the formation of necrotic granulomas. <u>A-C: Day 98 post CD infection (n=5 per group).</u> A) Representative histology images of lung sections. B) Principal component analysis of pathology scores. C) Mtb lung burden in Primary, BCG, and CoMtb Groups. <u>D-L: Day 35 post ULD infection (n= 10 primary, 5 CoMtb).</u> D) Representative confocal microscopy images demonstrating preserved alveolar integrity in non-necrotic Primary and CoMtb lesions. E) Representative confocal microscopy images depicting major cell populations within lesions. F) Percent of mice with necrotic lesions, covers two independent experiments. G) Heatmap showing cellular composition of clustered microenvironments. H) Representative map showing 50um^2^ neighborhoods, color-coded microenvironment. I) Percent area of lesion comprised by each microenvironment. Uninvolved regions (grey) not included. J) Percent of lesion comprised by necrotic region (pink). K) Ratio of lymphoid (blue, green) to myeloid (yellow, orange, pink) predominant regions. L) Relative density of PPD signal per 50 um^2^ neighborhood. Single-group comparisons by Mann-Whitney U test. *p < 0.05, **p < 0.01, ***p < 0.001, ****p < 0.0001. Error bars (F) reflect 95% confidence intervals. Points represent individual mice or lesions from individual mice. Data are representative of one (A-C) or two (D-L) independent experiments. See also Figure S1.

Detection of multiple distinct lesion types during primary infection, including both necrotic and non-necrotic lesions, raised the question of whether this reflected distinct stages of lesion progression (i.e., initial aerosol-seeded versus secondary, disseminated lesions) or an earlier divergence in lesion organization. To test these distinct possibilities, we utilized an ultra-low dose (ULD) aerosol infection (1-3 CFU), which results in the formation of a solitary organized lesion in most mice.^14^ Primary and CoMtb mice were assessed at d35 p.i., a timepoint shortly after the formation of mature lesions. In control animals, we observed formation of single lesions which possessed heterogeneous organization, with 15/27 mice across two experiments possessing granulomas containing a central necrotic core dominantly comprised of neutrophils (CD177-positive cells), necrotic debris (nuclear dye), and absence of alveolar epithelial staining (p120), consistent with destruction of the epithelial architecture (Fig 1D-F, S1C). In stark contrast, the remaining 12/27 lesions in control infected mice lacked this necrotic core and instead contained tightly aggregated clusters of antigen-bearing macrophages (CD68+, Siglec F-, PPD+) which were surrounded by intact alveolar epithelium (Fig 1D-F), consistent with alveolitis.

We next examined early lesions after ULD Mtb infection of mice with CoMtb infection. We observed complete absence of lesion necrosis, and instead these lesions again were comprised of tightly aggregated infected macrophages surrounded by intact alveolar epithelium, consistent with alveolitis. To quantify these findings, we used histo-cytometry and CytoMAP.^25,26^ We first segmented single cells to define major cell types within imaged tissues (neutrophils (CD177), macrophages (CD68), T cells (CD3, CD4), B cells (B220), Mtb antigen-bearing cells (PPD)), and also examined alveolar epithelial integrity with p120 staining (Fig S1D). We next used CytoMAP to raster-scan the spatial neighborhoods (radius = 50μm) within the imaging data and clustered these neighborhoods into discrete tissue region subtypes (i.e. microenvironments) based on the similarity of cellular composition. This analysis revealed that 4/10 sampled lesions in the setting of primary infection had regions consistent with necrosis (neutrophil enrichment, paucity of alveolar epithelium, R1/pink) and high antigen abundance (R2/orange), surrounded by regions of high myeloid density (R3/yellow) (Fig 1G-J). In contrast, lesions in CoMtb mice were highly enriched for lymphoid-dominant regions with intact p120 staining (R4/blue, R5/green), consistent with alveolitis (Fig 1I,K). Necrotic granulomas in control animals were also associated with increased PPD abundance as compared to non-necrotic lesions, while lesions in CoMtb mice had markedly reduced PPD abundance (Fig 1L), consistent with markedly decreased CFU at early timepoints (Fig 1C). Since most lesions in this ULD infection model at these early time points represent those seeded by the initial aerosol infection, this indicated that there is an early divergence in primary lesion development (necrosis vs. alveolitis) even in genetically identical mice infected with the same Mtb strain, and that concomitant immunity afforded by CoMtb abrogates formation of necrotic granulomas and leads to the generation of alveolitis.

### CoMtb alters the immune landscape following Mtb infection

We next sought to obtain a holistic understanding of lesion divergence at the early timepoints. For this, we performed spatial transcriptomics analysis using the Nanostring GeoMx platform on necrotic and non-necrotic lesions from primary ULD infected mice (d35 p.i.). We selected multiple regions of interest (ROI) within necrotic and non-necrotic lesions, as well as from uninvolved distal lung regions. ROI counts for each granuloma and uninvolved region were aggregated, normalized, and assessed by PCoA. This analysis revealed major transcriptomic distinctions between lesions and uninvolved tissues. Further, while ROIs from non-necrotic lesions were dispersed along PCoA 1 and overlapped with uninvolved tissues, ROIs from necrotic granulomas were tightly clustered and entirely distinct from uninvolved tissues (Fig 2A, S2A). Gene set enrichment analysis of comparing necrotic vs non-necrotic ROIs identified multiple pathways increased in necrotic ROIs, related to neutrophil biology and lesion necrosis, including type-I interferon production and signaling, neutrophil activation/trafficking (chemotaxis, phagocytosis, reactive oxygen, and nitrogen species production), cell death, TGFβ signaling, and tissue degradation/remodeling (Fig 2B). Together, this suggests that even in primary infection settings, individual lesions have vastly different immune and inflammatory landscapes, with a dominant difference being type I IFN and neutrophil-associated factors.

**Figure 2:**
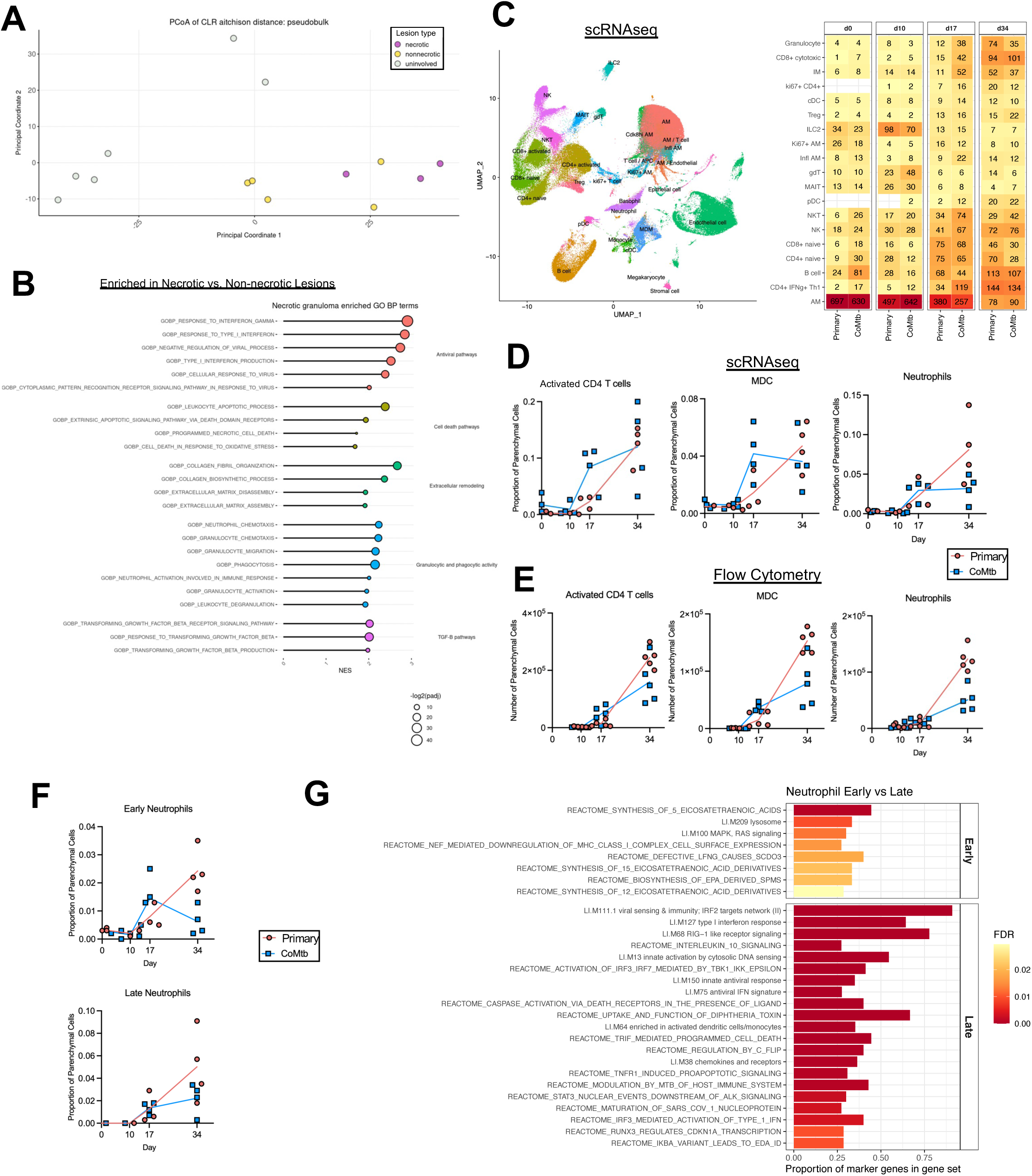
CoMtb alters the immune landscape following Mtb infection. <u>A-B: Day 35 post ULD infection, n = 8.</u> A) PCoA analysis of ROI transcriptomes, color coded by lesion type (necrotic vs non-nercotic, 3 ROIs per point) or location (uninvolved, 1 ROI per point)). B) GSEA analysis showing pathways enriched in primary necrotic lesions vs primary non-necrotic lesions. <u>C-E: Multiple timepoints post CD infection.</u> C) UMAP depicting cell types identified by scRNAseq analysis of lung parenchymal cells, heatmap showing changes in cellular abundance across timepoints, numbers reflect the median number of cells of a given type per thousand cells. D) Change in proportions of selected cell populations over time, as determined by scRNAseq. E) Change in numbers of selected cell populations over time, as determined by flow cytometry. F) Change in proportions of early and late neutrophil clusters over time, as determined by scRNAseq. G) GSEA analysis showing pathways enriched in early and late neutrophil clusters. Points represent individual lesions (A, 3 ROIs samples per lesion, 1 per uninvolved area), and individual mice (D, E, F). False discovery rate-adjusted p values determined using the R fgsea package. Data are representative of one (A-D, F-G) or two (E) independent experiments. See also Figures S2 and S3.

To gain further insights into lesion development and effects of CoMtb on immune responses during infection, we performed single-cell RNA sequencing (scRNAseq) and flow cytometry analysis of lungs from Mtb-infected animals with and without CoMtb immediately pre-infection and at 10, 17 and 34 days after CD infection. Clustering of scRNAseq data across timepoints and conditions allowed for robust identification of the major immune cell types comprising pulmonary lesions, including T cells, macrophages, neutrophils, and B cells (Figs 2C, S2B). Primary infection of mice resulted in gradual recruitment of T cells, monocyte-derived cells, as well as neutrophils which continued to accumulate over time (Fig 2C,D). In contrast, infection of CoMtb mice induced an increased representation of activated CD4 T cells (primarily defined by *CD44* and *IFNg*) and monocyte-derived cells (MDC) at early timepoints (d17), and this was correlated with decreased bacterial burdens at this timepoint (Fig 2D, S3A). These differences equalized by d34 (Fig 2D), when there was a greater increase in lung bacterial burdens in the primary group (Fig S3A). Neutrophils were found in equivalent representation at day 10 and 17 for both conditions, but continued to increase in control infected mice, and were closely correlated with bacterial burdens over time, while remaining stable in the CoMtb group (Fig 2D, S2C). Similar observations were confirmed by flow cytometry, demonstrating early increases in activated CD4 T cells and MDCs in CoMtb mice, and enhanced neutrophil abundance in primary Mtb settings at later time points (Figs 2E, S3B-D).

Additionally, we observed heterogeneity and distinct patterns of recruitment in different neutrophil populations by scRNAseq. We identified a cluster of neutrophils present within the lung prior to infection, “early neutrophils” which was enriched for pathways including eicosatetraenoic acids and T cell signaling and stimulation (Fig 2F,G, S2D). In CoMtb settings, this early neutrophil population had a more robust representation at d17 but declined by d34, and this contrasted the primary disease group where these cells continued to increase over the course of infection. A distinct “late” neutrophil cluster was identified at d17 p.i., and these cells showed a strong enrichment in signaling for type I IFN and cell death pathways. (Fig 2F,G, S2D). In primary infection, the late cluster neutrophils were markedly increased by d34, and this contrasted with CoMtb settings, which again demonstrated leveling off of neutrophil abundance at this timepoint (Fig 2F). Together this indicates that CoMtb-mediated pre-existing immunity results in pleiotropic effects on multiple innate and adaptive immune cell populations over the course of infection.

### CoMtb accelerates T cell and MDC activation, blunts neutrophil responses

To further dissect the transcriptional changes in distinct cell types in the presence or absence of CoMtb over time, we used GSEA. Even prior to aerosol infection, there were differences in pathways associated with cell cycle and mitochondrial respiration in ILC2s in the setting of CoMtb, but not primary infection, potentially indicating innate training, which is consistent with previous reports (Fig 3A).^27^ In agreement with our cellular abundance analysis, a stronger cell cycle/division response was seen in CD4 T cells following CoMtb at d17, indicating ongoing cellular activation and proliferation of T cells within the lung parenchyma. Starting d17 p.i., we also observed striking response differences in interferon response pathways which were upregulated over time across most cell populations (Fig 3A).

**Figure 3:**
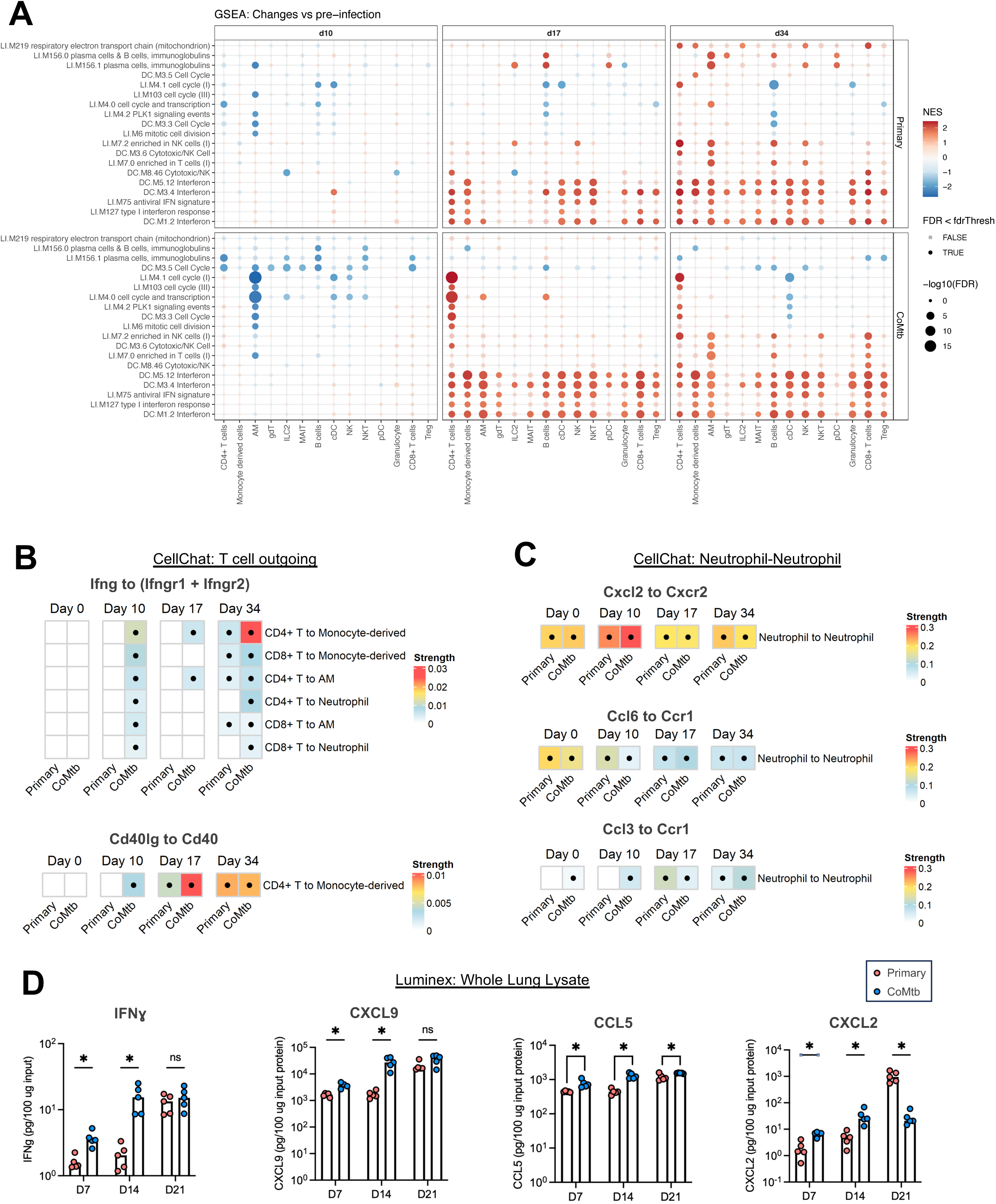
CoMtb accelerates T cell and MDC activation, blunts neutrophil responses. <u>Multiple timepoints post CD infection.</u> A) GSEA analysis showing pathways enriched following aerosol infection at days 10, 17, and 34 post infection, in the setting of primary infection and CoMtb. B) Predicted strength of selected T cell to myeloid cell signaling interactions quantified using CellChat. C) Predicted strength of significant neutrophil-neutrophil interactions in CellChat’s chemokine pathways. D) Levels of IFN□, CXCL9, CCL5, and CXCL2, measured by Luminex. False discovery rate-adjusted p values determined using the R fgsea package. Dots in B and C indicate strength is significantly higher compared to a null distribution (i.e., CellChat-reported p value < 0.05). Single-group comparisons in D by t test. Data are representative of one (A-C) or two independent experiments (D) experiments. See also Figures S2 and S3.

To elucidate what might be driving these interferon pathways, we used CellChat analysis^28^ to elucidate the predicted ligand-receptor interactions between T cells and myeloid cells. We identified markedly increased outgoing *Ifng* signals from CD4 and CD8 T cells and multiple myeloid subsets in the setting of CoMtb at very early time points (d10) (Fig 3B), suggesting enhanced activation of myeloid cells by T cells. This was supported by Luminex analysis of whole lung lysate, which demonstrated elevated levels of IFN□ and CXCL9 (downstream of IFN□) as early as d7 and d14 p.i. (Figs 3D, S3D). Similarly, outgoing *Cd40lg* interactions from T cells to *Cd40* on myeloid cells were increased d10 and d17 p.i. (Fig 3B). Given that these timepoints are prior to (d10), or shortly after the time (d17), when changes in bacterial burdens are observed (Fig S3A), and that both IFN□ and CD40L have important roles in Mtb immunity, these results suggest a causative role for T cell-derived activation of myeloid cells in CoMtb-mediated protection.

We also used CellChat to probe potential chemotactic interactions that could result in enhanced neutrophil recruitment, specifically focusing on neutrophil-neutrophil interactions that can drive feed-forward recruitment loops known to mediate tissue destruction in other models of inflammation.^29^ We found enhanced *Cxcl2-Cxcr2* interactions, and these were increased in CoMtb mice at early time points but were preferentially enriched in primary infected animals at later timepoints associated with enhanced neutrophil abundance (Fig 3C). Similarly divergent CXCL2 protein abundance between the groups was also confirmed via Luminex. We found modestly increased CXCL2 protein abundance in settings of CoMtb at early timepoints (d14), but this was then dwarfed by massive upregulation of CXCL2 in primary infected animals one week later (d21) (Fig 3D). Together, these data indicate that Mtb infection in CoMtb settings are associated with rapid recruitment and activation of T cells and enhanced IFN□ sensing by local myeloid cells, as well as with limited neutrophil recruitment at late timepoints. In contrast, primary infection in the absence of prior immunity is associated with limited early T cell activity and continued neutrophil influx over time.

### CoMtb shapes early tuberculous lesion cellularity and organization

The above data indicated that prior Mtb exposure induces a fundamental shift in the pulmonary immune landscape during early Mtb lesion formation which leads to divergent disease progression at later time points. To understand the organization of immune cells and investigate signaling microenvironments within developing lesions, we examined very early lesions 17 days p.i. using quantitative microscopy. In accord with our observations by scRNAseq and flow cytometry, we observed accelerated CD4 T cell responses in CoMtb infected mice, with a higher density of CD4 T cells within developing lesions, and particularly in neighborhoods in close proximity to PPD+ MDCs (Fig 4A, B). CoMtb lesions also demonstrated an increased proportion of MHCII+ MDCs, consistent with increased coupling of T cell activation with downstream myeloid cell maturation (Fig 4A, B).

**Figure 4:**
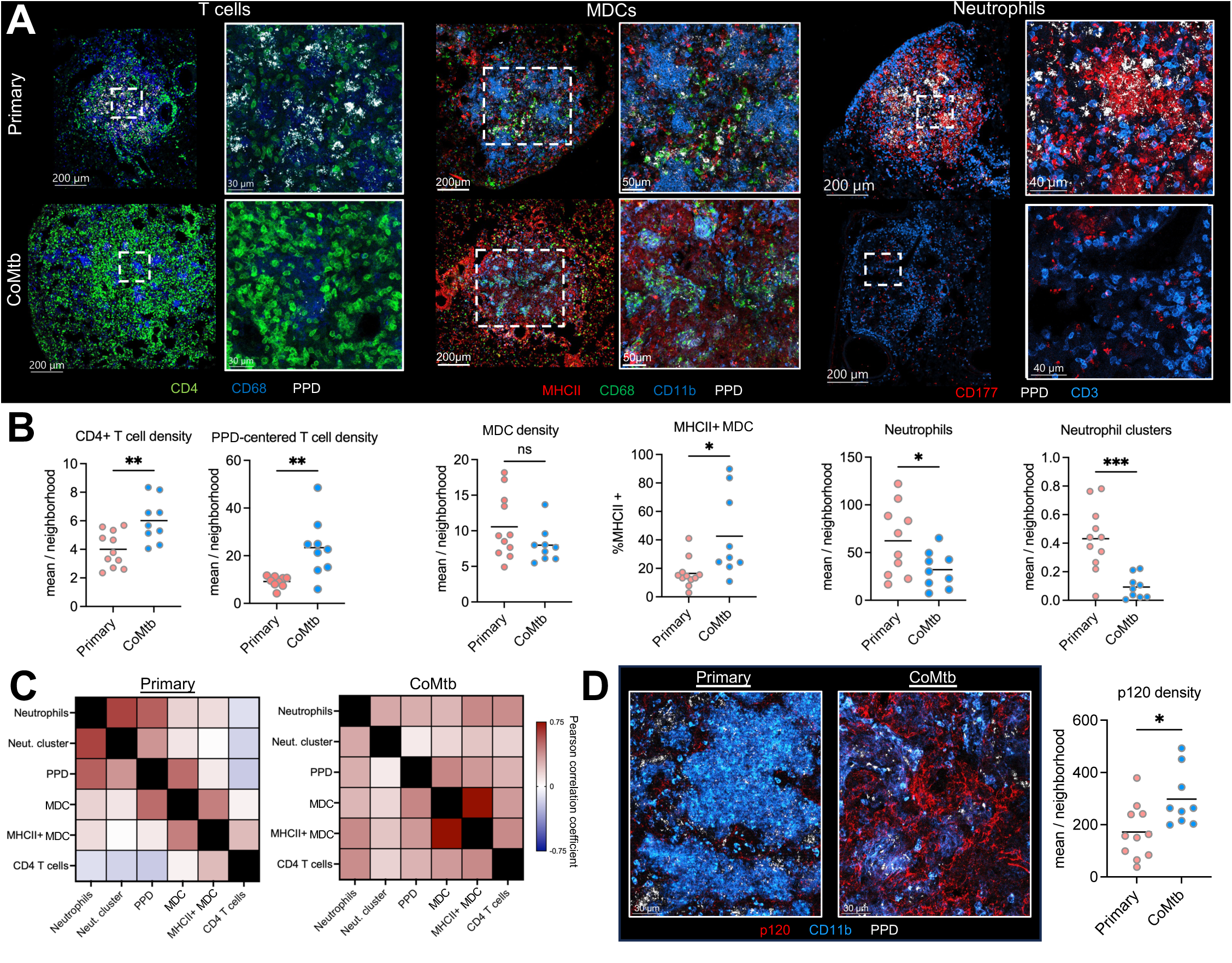
CoMtb shapes early tuberculous lesion cellularity and organization. <u>Day 17 post CD infection, n = 5 per group.</u> A) Representative confocal images showing lesions and zoom-ins highlighting T cells, MDCs, and neutrophils. B) Relative cellular density of these cell types within lesions as determined by histo-cytometry. C) Pearson correlation coefficients of the indicated cell populations within microenvironments. D) Confocal image and spots/neighborhood of p120 staining. Single-group comparisons in by unpaired t test. Correlations by Pearson’s correlation test. Points represent individual lesions. Data are representative of two independent experiments.

Many of the early lesions formed in the absence of prior immunity (primary) already possessed extensive neutrophil clusters, albeit we also observed extensive heterogeneity in this process, consistent with the divergent lesion outcomes seen at later timepoints (Fig 4A,B). In contrast, lesions formed in the setting of CoMtb possessed a much lower density of neutrophils and the infiltrating cells were sparsely distributed throughout the lesions (Fig 4A,B). The reduced neutrophil infiltration observed at this early timepoint by quantitative imaging differed markedly from the neutrophil cellularity observed at the same timepoint in flow cytometry and scRNAseq datasets which showed similar neutrophil cellularity across groups, potentially reflecting the inefficiency in recovering viable neutrophils in single cell suspensions especially when these cells are undergoing cell death, as seen in our spatial transcriptomics data (Fig 2B).^30^ To further understand the spatial relationships of different cell types with respect to one another, we analyzed the cell-cell correlations of cellular abundance across tissue neighborhoods within lesions. We found that even at this early time point, there were already distinct organizational features. In control mice without prior immunity, neutrophils were strongly associated with large clusters, as defined by aggregates >3000 μm^3^, which were also highly associated with local PPD antigen abundance (Fig 4C), and both were negatively correlated with T cells. These lesions also had signs of decreased alveolar integrity as compared to CoMtb, with a decreased density of p120+ cells (Fig 4D). In contrast, CoMtb generated lesions in which PPD antigen was positively correlated with both macrophages expressing MHCII and CD4 T cells, suggesting closer proximity and cross-talk between these cells. Together, this suggests that CoMtb has a dominant effect on shaping immune cell organization, abundance, and activation within early developing pulmonary lesions following aerosol Mtb infection.

### CD4 T cells are required for CoMtb-mediated protection from lesion necrosis

We hypothesized that the CoMtb-mediated acceleration of CD4 T cell responses was responsible for improving local myeloid responses and CFU burden. To test this, we depleted CD4 T cells in control and CoMtb infected mice using anti-CD4 antibody, with the depletion beginning one day prior to infection, and examined lesion structures and lung CFU 35 days later (Figs 5A, S4A). In stark contrast to aerosol Mtb-challenged CoMtb mice which completely lacked necrotic lesions, CD4 T cell-depleted CoMtb mice developed highly necrotic lesions which contained a central core that was densely packed with infiltrating neutrophils and which lacked epithelial staining (Figs 5B, 5D-F, S4B). Depletion of CD4 T cells also led to a near-complete reversion of bacterial protection offered by CoMtb, resulting in minimal differences in lung CFU between control and CoMtb infected CD4-depleted mice (Fig 5C). Together, this suggests that CD4 T cells play a pivotal role in regulating neutrophil abundance and lesion necrosis and are a major contributing factor mediating the CoMtb-reduction in bacterial burdens.

**Figure 5:**
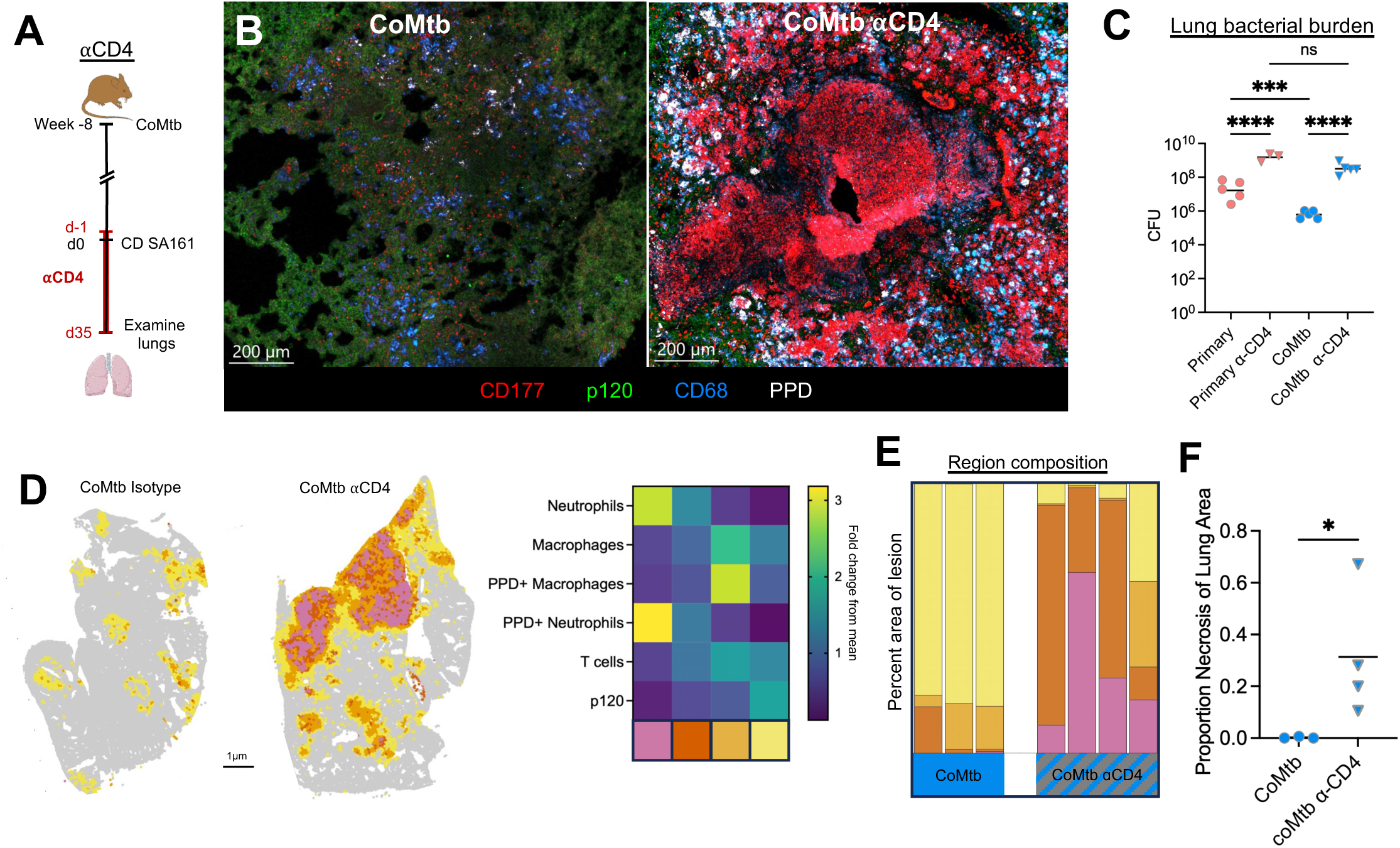
CD4 T cells are required for CoMtb-mediated protection from lesion necrosis. <u>A) Experimental outline. Subset of mice received CoMtb, then all mice aerosol infected with CD Mtb. Mice then received □CD4 depleting antibody or isotype from d-1 until harvest. n = 3-5 per group.</u> B) Representative confocal images showing presence of necrosis with □CD4 administration. C) Pulmonary bacterial burdens. D) Representative map showing 50um^2^ neighborhoods, color-coded microenvironment and heatmap showing cellular composition of clustered microenvironments. E) Percent area of lesion comprised by each microenvironment. Uninvolved regions (grey) not included. F) Percent of lesion comprised by necrotic region (pink). Single-group comparisons by Mann-Whitney U test. Points represent individual mice. Data are representative of two independent experiments. See also Figure S4.

### Neutrophils drive lesion necrosis

Given the correlation in neutrophil abundance, increased bacterial burdens, and severe pathology that we and others have observed, we next hypothesized that neutrophils were necessary for pulmonary lesion necrosis and promote enhanced bacterial replication. The role of neutrophils during Mtb infection is multifaceted, with evidence suggesting both beneficial roles for bacterial control, and detrimental roles driving worsened outcomes, especially in severe disease. Neutrophil depletion has been shown to compromise control of Mtb bacterial burden in the C3H model, though the impact of neutrophils on determining lesion organization has not been examined directly.^31–34^ To directly test the role of neutrophils in promoting granuloma formation in the C3H mouse model, we infected mice with a CD of Mtb, and depleted neutrophils with an anti-Ly6G (□Ly6G, IA8) antibody starting at d7 p.i., when Mtb first starts to infect non-alveolar macrophage cell types, to d28 p.i., when mature necrotic lesions have formed (Fig 6A, S5A). Neutrophil depletion resulted in a complete abrogation in lesion necrosis (Fig 6B), instead generating lesions comprised of alveolitis, akin to CoMtb, with less overall extent of lung involvement (Fig 6D-F). Neutrophil depletion also resulted in an approximately 2-log reduction in lung bacterial burdens (Fig 6C), together indicating that neutrophils promote lesion necrosis and restrict Mtb immune control.

**Figure 6:**
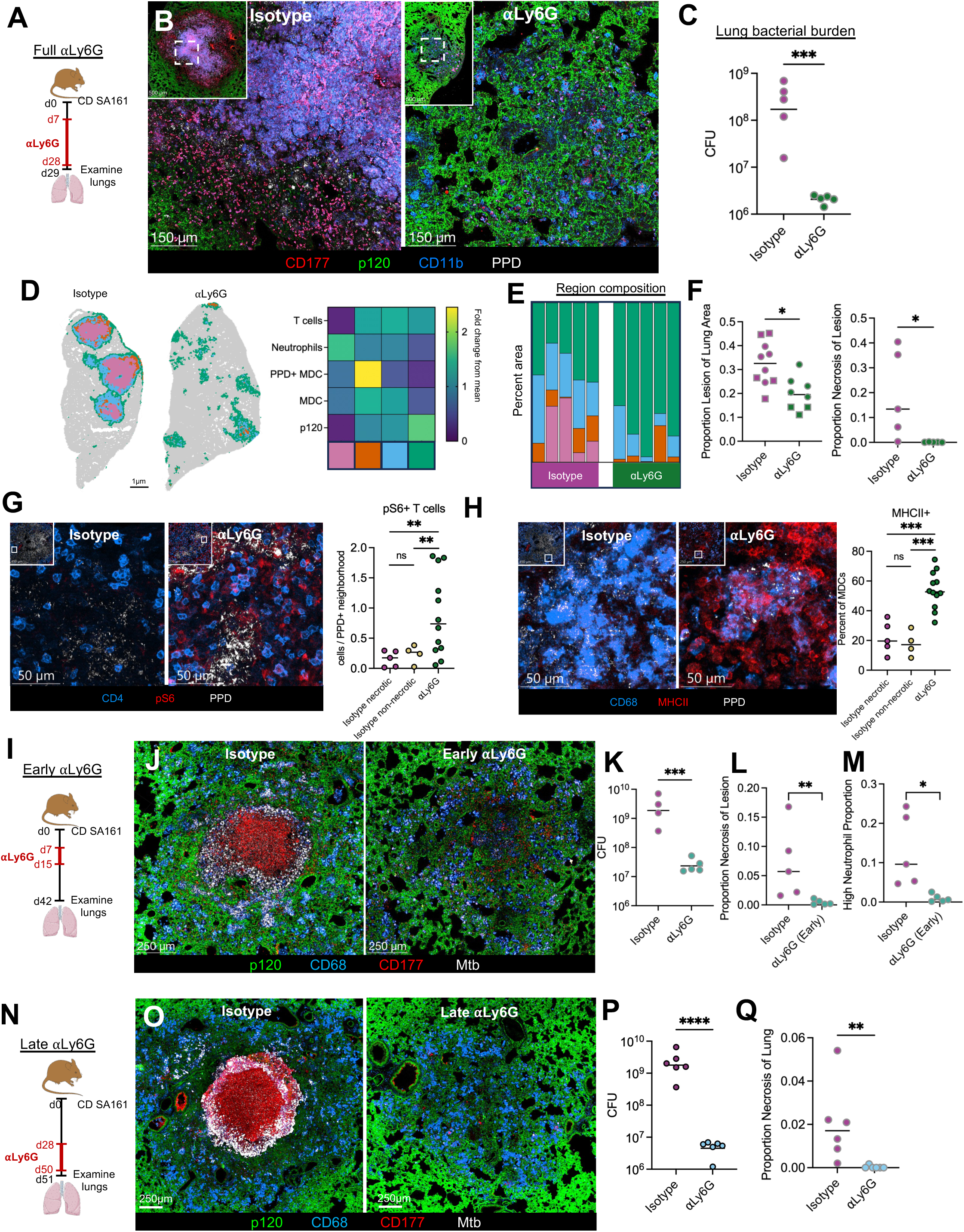
Neutrophils drive lesion necrosis. A) Experimental outline for A-H. Mice received CD aerosol infection, then administered □Ly6G depleting antibody or isotype from d7-d28, lungs taken d29. B) Representative confocal images showing abrogation of necrosis with □Ly6G administration. C) Pulmonary bacterial burdens. D) Representative map showing 50um^2^ neighborhoods, color-coded microenvironment and heatmap showing cellular composition of clustered microenvironments. E) Percent area of lesion comprised by each microenvironment. Uninvolved regions (grey) not included. F) Percent lesion (any color) of total lung area, and percent of lesion comprised by necrotic region (pink). G) Representative confocal images and quantification showing increased pS6+ T cells following □Ly6G administration. H) Representative confocal images and quantification showing increased MHCII+ in MDCs following □Ly6G administration. I) Experimental outline for I-M. Mice received CD aerosol infection, then administered □Ly6G depleting antibody or isotype from d7-d15, lungs taken d43. J) Representative confocal images showing abrogation of necrosis with “Early” □Ly6G administration. K) Pulmonary bacterial burdens. L) Percent of lesion comprised by necrotic region (pink). M) Percent of lesion comprised by microenvironments with high neutrophil density. N) Experimental outline for N-Q. Mice received CD aerosol infection, then administered □Ly6G depleting antibody or isotype from d28-d49, lungs taken d50. O) Representative confocal images showing decreased necrosis with “Late” □Ly6G administration. P) Pulmonary bacterial burdens. Q) Percent of lung area comprised by necrotic region (pink). Single-group comparisons by unpaired t test (C, F left, K, P) or Mann-Whitney U test (F right, G, H, L, M, Q). Points represent individual mice. Data are each representative of three (A-H) or two (I-Q) independent experiments. See also Figure S5.

To explore how neutrophils affect local immune landscapes within lesions, we again performed quantitative image analysis. Lesions in neutrophil-depleted mice exhibited enhanced infiltration of CD4 T cells into the Mtb-infected, macrophage-rich, central granuloma cores, including CD4 T cells with increased pS6 staining, suggesting recent TCR signaling, directly adjacent to PPD+ myeloid cells (Fig 6G). We also observed markedly increased MHC-II staining in neighboring cells, likely reflecting local inflammatory signaling and myeloid cell activation (Fig 6H).

We next hypothesized that neutrophil recruitment during the early stages of lesion formation might shape the downstream events of lesion progression. To test this, we administered □Ly6G depleting antibody 7-15 days p.i. (early depletion), then waited an additional 4 weeks to allow for lesion development (Fig 6I). When we evaluated the lungs of these mice at d42, we observed that early neutrophil depletion completely blocked lesion necrosis, instead driving generation of alveolitis. Early neutrophil depletion also resulted in a 2-log decrease in lung bacterial burdens, comparable to that observed using the extended depletion protocol (Fig 6J-L, S5B). Neutrophils were still observed in these lesions, albeit at lower numbers as compared to full-depleted animals, but the cells that did infiltrate did not exhibit extensive clustering (Fig 6M). Together this suggests that neutrophil influx during the very initial stages of lesion formation shape granuloma development and downstream disease progression.

Finally, we examined whether continued neutrophil recruitment at later stages of infection, after the necrotic lesions have been established, is required for maintenance of disease pathology. To test this, we administered □Ly6G antibody 28 days after CD infection with SA161 Mtb, and continued treatment for two weeks (late depletion) (Fig 6N). Evaluation of these lungs via imaging revealed that late neutrophil depletion also had marked beneficial effects on tissue pathology. Nearly all lesions in late depleted animals lacked necrosis and caseation, and we found only a single lesion in 1/6 mice containing a small necrotic center (Figs 6O,Q, S5C,D). Further, late neutrophil depletion was associated with a 2.5 log reduction in lung bacterial burdens (Fig 6P). Together, this suggests that continued neutrophil recruitment at late timepoints is required to propagate lesion necrosis and restrict immunity against Mtb.

## Discussion

From the earliest pathologic examinations of TB, it has been apparent that Mtb infection results in pulmonary lesions with vastly different organization, which range from generation of necrotizing and cavitating lesions to pneumonia-like alveolitis, and these are known to have major implications for disease severity and resiliency to antibiotic therapy.^35^ However, dissecting the mechanistic basis of these divergent processes in humans has been challenging due to extensive genetic and environmental variation among the populations, differences in past exposure history, including BCG immunization or infection with either Mtb or nontuberculous mycobacteria, and based on antibiotic usage. Some historical studies examined differences between Mtb lesion structures in vaccinated versus unvaccinated animals, but lacked the modern tools to examine cellular organization and interactions in a quantitative manner. More recent studies have focused on granuloma structures in animals in the absence of prior immunity. To investigate how pre-existing immunity affects lesion composition and organization we used a mouse model for concomitant immunity (CoMtb) in C3H mice. We find that prior immunity rapidly reshapes lesion pathology by abrogating the formation of necrotic granulomas and instead leading to generation of alveolitis with reduced bacterial burdens, and that these changes are dominantly dictated by the opposing roles served by CD4 T cells and neutrophils. Together, this study uncovers major cellular mechanisms leading to the divergence in lesion structure and disease progression based on immune history and lends insights into the mechanisms underlying the differences in pathology caused by primary and post-primary TB.

Akin to our previous work examining responses to Mtb infection following BCG immunization,^36^ we show that the pre-existing immunity conferred by CoMtb accelerates the localization of T cells and monocyte-derived cells to lesions and enhances their activation state at the site of infection, and this in turn is associated with a blunted neutrophil response at late timepoints. This results in a complete abrogation of lesion necrosis, which we demonstrate is CD4 T cell-dependent.^37^ While an important role for CD4 T cells in controlling Mtb infection has been appreciated for decades, our study demonstrates how this immunity is achieved at the tissue level and in the context of lesion composition and architecture. In settings without prior exposure, the activation of T cells and their recruitment to infected sites within the lung is delayed and occurs only after the early onslaught of infiltrating innate cells, including neutrophils. This allows for the establishment of early pro-necrotic lesions which we see developing as early as d17 p.i., and these early lesions likely already have the ability to suppress local adaptive immune responses, promote further neutrophil recruitment, and provide a safe harbor within the developing core for enhanced bacterial replication. In contrast, pre-existing immunity elicits rapid recruitment of T cells to the infected sites and these in turn promote local monocyte and macrophage activation, together restraining the establishment of pro-necrotic lesion centers and unchecked bacterial replication. Additional differences between primary TB and settings of prior immunity may also arise from training of innate responses, including that of alveolar macrophages, monocytes, and neutrophils, and these could directly impact responses to Mtb or indirectly alter crosstalk with CD4 T cells.^27,38^

Potential mechanisms for the protection afforded by CD4 T cells as elucidated by our scRNAseq analysis include increased IFN□ production by T cells and local sensing by myeloid cells, which would promote enhanced bacterial control during the earliest stages of infection. Moreover, IFN□ signaling in non-hematopoietic stromal cells as well as intrinsically in neutrophils has been shown to reduce neutrophil localization to the Mtb-infected lung, suggesting pleiotropic effects.^39,40^ Additional candidate mechanisms include differential activation of monocyte-derived cells via CD40L or MIF, which are both important for optimal bacterial control,^41,42^ as well as regulation of myeloid trafficking via CD6-ALCAM interactions.^43^ Of note, our findings that every lesion underwent necrosis in the setting of CD4 depletion is distinct from what we observe during ULD infection of C3H mice in the absence of prior immunity, in which only a subset of lesions develop necrosis. Thus, our data show that CD4 T cells are essential for preventing lesion necrosis in settings of prior immunity, suggesting that their rapid recruitment and function within early developing lesions dominantly shapes downstream disease progression.

While pre-existing immunity may be one manner which promotes alveolitis following Mtb infection, there is evidence that several other factors may also influence lesion structure. For example, the lesion heterogeneity that we observed in isogenic mice infected with genetically identical bacteria suggest that stochastic events, such as differences in the activation phenotype of the first cell to uptake Mtb or early cellular interactions in distinct regions of the lung, may also lead to necrotizing granulomas in some cases and alveolitis in others. Mtb strain characteristics and host genetics also likely contribute, as large necrotic granulomas form more readily in C3H mice infected with the Mtb SA161 strain than with the H37Rv strain,^15^ whereas infection by either Mtb strain in C57BL/6 mice, which mount a robust Th1 response, leads to lesions comprised of alveolitis even in the absence of prior immunity.^44^ Further evidence that genetic differences in the Mtb strains themselves can drive different lesion types comes from recent work showing that clinical Mtb strains associated with high transmission in human populations induce more granuloma necrosis in C3H mice than Mtb strains associated with low transmission. Environmental factors, including co-infections, may also shape Mtb lesion organization. Clinical studies demonstrate that HIV infected patients with low CD4 T cell counts are less likely to form cavities.^45^ Initially, this seems incongruent with our findings that the absence of CD4 T cells strongly promotes necrosis. However, the CD4 depletion in HIV-infected individuals does not occur in isolation. These individuals are viremic, often have additional co-infections, and their immune systems are globally dysregulated, all of which may influence lesion structure.^46^ Thus, while pre-existing immunity strongly influences lesion structure, additional work is needed to dissect the mechanisms driving lesion progression and disease pathogenesis across different settings and clinical scenarios.

Neutrophils have a strong association with severe disease in tuberculosis, as shown in several mouse models, experimentally and using computational modeling in NHPs, and observationally in clinical studies.^47–50^ This has led to the proposal of a “tipping point” model, where neutrophils mediate disease exacerbation downstream of multiple mechanisms of impaired host resistance.^51^ We now further this model showing that neutrophils actively regulate lesion organization and negatively affect disease pathology, and that they are required for this throughout the different phases of infection. We find that neutrophils are essential very early during infection to drive necrosis and enhance disease severity, and even when neutrophils are later given the chance to enter tissues, they do not display the same magnitude of recruitment, do not drive necrosis, nor markedly affect bacterial burdens. This suggests that there is a brief window for neutrophils to cause lesion necrosis, and if neutrophils are not present during that time, responses by other cell types such as CD4 T cells or macrophages dominantly shape local tissue environments that alter downstream disease progression. Mechanistically, it is likely that in addition to their role in driving necrosis, early clusters/swarms of recruited neutrophils locally impair immune responses, and we see reduced T cell localization and TCR sensing near antigen-bearing cells, and this is associated with less downstream myeloid cell activation. In addition to early time points, we find that neutrophils are required for sustaining disease pathology even after initial necrotic lesion formation, and that depletion of neutrophils after formation of necrosis leads to dramatic improvements in lesion pathology and marked reduction in lung bacterial burdens. To our knowledge, this represents the largest improvement seen in pulmonary bacterial burdens with host-directed therapy initiated at late timepoints. This also represents a more clinically relevant scenario, since patients present to clinic late after exposure and almost always with radiographically-apparent pulmonary lesions.^52,53^ Thus, our work builds on the “tipping point” model by elucidating the role of neutrophils in shaping lesion structure, as well as demonstrate that neutrophils do not simply respond to immune failure at chronic timepoints, but act during early inflection points to drive downstream disease progression. The potential mechanisms by which neutrophils mediate this process include NETosis^15,18^ driving type I IFN production,^18^ and ROS,^31^ warranting careful evaluation in future studies.

Overall, our work establishes a mouse model to dissect how pre-existing or concomitant immunity modifies disease progression and leads to the formation of distinct lesion types. Clinically, necrotic pulmonary lesions pose a significant challenge for antibiotic treatment, in large part due to the reduced penetration of antibiotics into necrotic centers. Necrotic lesions that have emptied their caseous core to form cavitary lesions also pose an increased risk of relapse and long-term pulmonary sequelae, such as impaired clearance of respiratory secretions and recurrent infections.^11,12^ Our demonstration that neutrophil depletion, even administered after granuloma formation, can limit lung destruction and preserve alveolar epithelium architecture, suggests that neutrophils may provide a useful target for host-directed therapy in conjunction with antibiotic treatment. While an indiscriminate neutrophil depletion is not a practical clinical solution due to the overwhelming risk of other infections, multiple inhibitors of neutrophil trafficking and activation are currently developed for other indications, and may be useful in reducing detrimental pathology and potentially even shortening treatment courses.

## Acknowledgements

This study was supported by NIH contract 75N93019C00070 (K.B.U., M.Y.G) and NIH grants 5K08AI166072 (B.H.G.) and 1U19AI162583 (K.B.U., B.H.G.).

## Contributions

Conceptualization, B.H.G., M.Y.G., and K.B.U.; Formal analysis, B.H.G., J.M.K., K.A.F., S.B.C., C.R.P., F.J.D., M.L.N., M.H., A.T.G., A.H.D., M.G., J.D.A., M.YG., K.B.U.; Funding acquisition, B.H.G., M.Y.G., and K.B.U.; Investigation, B.H.G., J.M.K., K.A.F., S.B.C., C.R.P., F.J.D., M.L.N., M.H., A.T.G., A.H.D., A.A., M.G., J.D.A., M.Y.G., and K.B.U.; Methodology, B.H.G., M.Y.G., J.D.A., M.Y.G., and K.B.U.; Project administration, B.H.G., M.Y.G., and K.B.U.; Resources, B.H.G., A.A., M.Y.G., J.D.A., M.Y.G., and K.B.U.; Supervision, B.H.G., M.Y.G., and K.B.U.; Validation, B.H.G., M.Y.G., and K.B.U.; Visualization, B.H.G., J.M.K., K.A.F., S.B.C., C.R.P., F.J.D., M.L.N., M.H., A.T.G., M.Y.G., and K.B.U.; Writing – original draft, B.H.G., M.Y.G., and K.B.U.; Writing – review & editing, B.G., J.M.K., C.R.P., F.J.D., M.L.N., M.Y.G., and K.B.U.

## STAR Methods

### Resource availability

#### Lead contact

Further information and requests for resources and reagents should be directed to and will be fulfilled by the Lead Contact, Kevin Urdahl.

#### Materials availability

This study did not generate new unique reagents.

#### Data and code availability

The mouse lung scRNAseq and spatial transcriptomics data generated during this study will be made publicly available upon publication.

## Experimental model and subject details

### Mice

C57BL/6 and C3HeB/FeJ mice were purchased from Jackson Laboratories (Bar Harbor, ME). All mice were housed in individually ventilated cages in specific pathogen-free conditions (maximum 5 mice/cage) within rooms with negative pressure ventilation and air filtering at Seattle Children’s Research Institute (SCRI). Animals were monitored under care of full-time staff, given free access to food and water and maintained under 12-hour light and dark cycles, with temperature controlled between 22-25 degrees Celsius. All possessed normal health and immune status. None had previous treatments, procedures, nor invasive testing prior to the initiation of our studies. Experiments were performed in compliance with the SCRI Animal Care and Use Committee. All experiments were conducted with sex and age-matched mice (both male and female mice between the ages of 8-12 weeks). The influence of sex was not assessed.

### Mycobacterium tuberculosis (Mtb)

For use in murine infections, Mtb SA161 strain was provided by Ian Orme (Colorado State University).^54^

## Method details

### Aerosol infections

Infections were done with a stock of Mtb SA161, as described previously. ^55^To perform CD aerosol infections, mice were placed in a Glas-Col aerosol infection chamber, and 50-100 CFU were deposited into their lungs. To confirm the infectious inoculum, two mice per infection were euthanized on the same day of infection, then their lungs homogenized and plated onto 7H10 or 7H11 plates for determination of CFU. To perform ULD aerosol infections, mice were placed in a Glas-Col aerosol infection chamber, and 1-3 CFU were deposited into their lungs.^14^

### CFU determination

Mouse organs (such as right or left lung, spleen) were individually homogenized in an M tube (Miltenyi) containing BS+0.05% Tween-80. The resulting homogenates were diluted and plated onto 7H10 plates. Plates were incubated at 37 degrees Celsius for a minimum of 21 days before CFU enumeration.

### Concomitant Mtb model (CoMtb)

The CoMtb model was established as described previously.^23,24^ Briefly, mice were first anesthetized by intraperitoneal injection of 400 ul of ketamine (4.5 mg/ml) and xylazine (0.5 mg/ml) diluted in PBS. Mice were placed in a lateral recumbent position, and the ear pinna was flattened with forceps and pinned onto an elevated dissection board using a 22 G needle. H37Rv Mtb grown to an OD between 0.2-0.5 over a 48-hour period was diluted to 10^6^ CFU/ml in PBS, and 10 ul (10^4^ CFU) was administered into the dermis of the ear using a 26s G Hamilton syringe. Mice were then rested for 6-8 weeks prior to subsequent aerosol challenge.

### Antibody depletions

For CD4 depletion studies, 500μg of an anti-CD4 depleting antibody (clone GK1.5) was administered intraperitoneally to mice once weekly, from the day prior to aerosol infection until harvest. For neutrophil depletion studies, 200ug of an anti-Ly6G depleting antibody (clone IA8) was administered intraperitoneally to mice three times weekly for the specified timepoints.

### Histology

Lungs processed for histology were fixed in 10% formalin for 24 hours, then dehydrated in 70% ethanol at 4 degrees for at least 24 hours. Samples were paraffin embedded and sectioned at the University of Washington Histology Core. Subsequently, slides were reviewed by a veterinary pathologist and scored in a blinded fashion based on the following metrics (see table S1): mixed granulomas (Ill-formed granulomas with mixture of macrophages and lymphocytes), defined granulomas (Well defined with increased separation of macrophages, epithelioid or multinucleated giant cells (MNGC) with lymphoid aggregates), perivascular lymphoid aggregates (PV LA), peribronchiolar lymphoid aggregates (PB LA), histiocytes, foamy macrophages, multinucleated giant cells, alveolar hyperplasia, neutrophils, necrosis, cholesterol clefts, edema, extent 1 (percent involvement of the lung), extent 2 (percent involvement of the lung in the worst manner).

### Lung single cell suspensions

At the indicated times post-infection, mice were anesthetized with isoflurane and administered 1 ug anti-CD45.2 antibody intravenously. After 5-10 minutes of in vivo incubation, mice were euthanized by CO_2_ asphyxiation. Mouse lungs were excised and lightly homogenized in HEPES buffer containing Liberase Blendzyme 3 (70 μg/ml; Roche) and DNaseI (30 μg/ml; Sigma-Aldrich) using a gentleMacs dissociator (Miltenyi Biotec). The lungs were then incubated for 30 min at 37°C and then further homogenized a second time with the gentleMacs. The homogenates were filtered through a 70 μm cell strainer, pelleted for RBC lysis with RBC lysing buffer (Thermo), and resuspended in FACS buffer (PBS containing 2.5% FBS and 0.1% NaN_3_).

### Antibody staining

Single cell suspensions were first washed in PBS and then incubated with 50 μl Zombie UV viability dye (BioLegend) for 10 min at room temperature in the dark. Viability dye was immediately quenched by the addition of 100 μl of a surface antibody cocktail diluted in 50% FACS buffer/50% 24G2 Fc block buffer using saturating levels of antibodies. Surface staining was performed for 20 min at 4°C. Then, the cells were washed once with FACS buffer and fixed overnight with the eBioscience Intracellular Fixation and Permeabilization kit (Thermo Fisher). The following day, cells were permeabilized with the provided permeabilization buffer, incubated for 20 min at 4°C with 100 μl of an intracellular antibody cocktail diluted 1:100 in permeabilization buffer, and washed with FACS buffer. Cells were analyzed on a BD Symphony A5 cytometer (BD).

### Antibodies

The following antibodies were used for staining mouse tissue sections for imaging or isolated cells for flow cytometry: B220 PCPCy5.5 (clone RA3-6B2; Biolegend), B220 PE/Fire 700 (clone RA3-6B2; Biolegend), CD103 PE (clone 2E7; Biolegend), CD105 R718 (clone MJ7/18; BD), CD11b BV480 (clone M1/70; BD), CD11b BV570 (clone M1/70; Biolegend), CD11b PCPCy5.5 (clone M1/70; Biolegend), CD11b PE/Fire 640 (clone M1/70; Biolegend), CD11b R718 (clone M1/70; BD), CD11c BV480 (clone HL3; BD), CD11c BV711 (clone N418; Biolegend), CD11c PCPCy5.5 (clone N418; Biolegend), CD11c PE (clone HL3; BD), CD177 AF647 (clone Y127; BD), CD177 CF555 [conjugated in house] (clone Y127; BD), CD177 CF633 [conjugated in house] (clone Y127; BD), CD177 PE (clone 1171A; R&D Systems), CD19 PE/Dazzle 594 (clone 6D5; Biolegend), CD26 PE-Cy7 (clone H194-112; Biolegend), CD3 BV480 (clone 17A2; BD), CD3 BV785 (clone 17A2; Biolegend), CD3 CF633 [conjugated in house] (clone 17A2; Biolegend), CD3 PE/Fire 640 (clone 17A2; Biolegend), CD3 PE/Fire 700 (clone 17A2; Biolegend), CD3e BUV737 (clone 145-2C11; BD), CD4 BV510 (clone RM4-5; Biolegend), CD4 CF594 [conjugated in house] (clone RM4-5; Biolegend), CD4 PE/Fire 700 (clone GK1.5; Biolegend), CD44 BV711 (clone IM7; BD), CD45.2 AF700 (clone 104; Biolegend), CD45.2 APC (clone 104; Thermo Fisher), CD45.2 R718 (clone 104; BD), CD62L AF488 (clone MEL-14; Biolegend), CD64 PerCP-eF 710 (clone X54-5/7.1; Thermo Fisher), CD68 BV421 (clone FA/11; BD), CD68 CF514 [conjugated in house] (clone FA-11; Thermo Fisher), CD68 CF750 [conjugated in house] (clone FA-11; Thermo Fisher), CD69 PE/Dazzle 594 (clone H1.2F3; Biolegend), CD86 BUV737 (clone 2331 (FUN-1); BD), CD8a BUV661 (clone 53-6.7; BD), Col1A1 CF660c [conjugated in house] (clone E8F4L; Cell Signaling), Col1A1 CF750 [conjugated in house] (clone E8F4L; Cell Signaling), CTLA4 (CD152) BV421 (clone UC10-4B9; Biolegend), CXCL2 CF555 [conjugated in house] (polyclonal; R&D Systems), FoxP3 AF700 (clone FJK-16s; Thermo Fisher), Gamma Delta TCR BUV805 (clone GL3; BD), iNOS AF405 (clone C-11; Santa Cruz Biotechnology), iNOS CF633 [conjugated in house] (clone CXNFT; Thermo Fisher), Ki67 BV605 (clone 16A8; Biolegend), Ki67 BV650 (clone 11F6; Biolegend), Ki67 eF506 (clone SolA15; Thermo Fisher), KLRG1 BUV395 (clone 2F1; BD), Ly6C AF700 (clone HK1.4; Biolegend), Ly6G BV605 (clone 1A8; Biolegend), MHCII (I-Ab) FITC (clone KH74; Biolegend), MHCII (I-Ak) FITC (clone 10-3.6; Biolegend), MHCII AF700 (clone M5/114.15.2; Biolegend), MHCII BV480 (clone M5/114.15.2; BD), Mtb FITC (polyclonal; Abcam), NOS2 APC eF780 (clone CXNFT; Thermo Fisher), p120 AF488 (clone 6H11; Santa Cruz Biotechnology), p120 AF594 (clone 6H11; Santa Cruz Biotechnology), Phospho-S6 CF750 [conjugated in house] (clone 2F9; Cell Signaling), Siglec F BV421 (clone E50-2440; BD), Siglec F BV480 (clone E50-2440; BD), SIRPα BV421 (clone P84; BD), T-bet PE-Cy7 (clone 4B10; Biolegend).

### Confocal microscopy

Lungs were removed and placed in BD Cytofix diluted 1:3 with PBS for 24hr at 4°C. Lungs were then washed two times in PBS and incubated in 30% sucrose for 24 hours at 4°C. Lungs were then embedded in OCT and freezing in a dry ice slurry with 100% ethanol. A CM1950 cryostat (Leica) was used to generate 20μm sections. Sections were rehydrated with 0.1M TRIS for 10 minutes, incubated for 1 hour at room temperature with blocking buffer (0.1M TRIS with 1% normal mouse serum, 1% bovine serum albumin, and 0.3% Triton X100), and then stained for 6 hours to overnight at room temperature with fluorescently conjugated antibodies. Following staining, slides were washed with 0.1M TRIS for 30 minutes and subsequently cover-slipped with Fluoromount G mounting media (SouthernBiotech). Images were acquired on a Leica Stellaris8 confocal microscope. For visual clarity, thresholds were applied to the displayed channel intensities in Imaris with identical settings applied across experimental groups.

### Histo-cytometry

Histo-cytometry analysis was performed as described previously, with only minor modifications.^26^ First, multiparameter confocal images were corrected for fluorophore spillover. Single color controls were made by mixing fluorophore-conjugated antibodies with Fluoromount G mounting media (SouthernBiotech) on a slide, then cover-slipping and collecting images with the same settings used for tissue imaging. Next fluorophore spillover was calculated and corrected using the Channel Dye Separation module in LAS X (Leica). Cell surfaces were created using Nucspot 750/780 nuclear staining using the Imaris surface creation module. Surfaces around neutrophils clusters were created on CD177 signal using the Imaris surface creation module (without splitting) followed by the application of size exclusion to only include surfaces >300 μm^3^. The location of PPD and p120 signal was determined using the Imaris spot creation module. The surface object and spot statistics were exported as CSV files. Object statistics were concatenated into CSV files and imported into FlowJo software for hierarchical gating.

### CytoMAP Spatial Organization Analysis

Spatial organization analysis was performed using CytoMAP.^25^ In brief, the position of all cell objects within tissues was used for virtual raster scanning with 50-µm radius neighborhoods. Raster-scanned neighborhoods were also used for clustering based on cell type abundance (cell types used denoted in associate heat maps) to identify distinct region types, and these regions were used for heatmap and positional visualization of regions. For figure 4B, cell centered neighborhoods with 50-µm radius were created around PPD+ cells, and the T cell density within these regions was calculated. The Pearson correlation coefficient was calculated for the number of cells of the different cell types within these neighborhoods.

### GeoMx DSP

A CM1950 cryostat (Leica) was used to generate 10μm sections from lungs processed as outlined above, then stored at −80°C. During the sectioning process, lesions were classified as necrotic or non-necrotic by visual inspection (presence of caseum) and brightfield microscopy (assessing alveolar integrity and presence of necrotic debris). The fixed frozen sample slides were baked for 2 hours at 60°C to ensure lung tissue adhered to slides.

Following baking, we performed target retrieval for 20 minutes following all recommended settings (MAN-10115-04). RNA targets were exposed using recommended concentration and duration of proteinase K (1ug/ml for 15 min). *In situ* probe hybridization took place overnight (18 hours) using standard hybridization solution with no custom spike-in (v1.0) Mouse NGS Whole Transcriptome Atlas RNA - lot # MWTA12002). The next day, off target probes were removed using stringent washes as recommended. Finally, morphology markers (SYTO13, B220 – PE, CD3 – CF594, and CD11b eF660) were added. Following antibody staining, slides and collection plate were loaded into GeoMx DSP instrument as recommended (MAN-10152-01). Slides were identified and records created for each.

Scan parameters were set for each channel: FITC/525 was utilized for SYTO13 nuclear staining, with exposure time of 50ms. Cy3/568nm was used for Alexa 532 to detect B220. Texas Red/615nm was used for Alexa 594 to detect CD3. Cy5/666nm was used for Cy5 to detect CD11b. All non-nuclear exposures were set for 200ms. Configuration files were obtained from the nanostring website. Syto 13 was used for focus. Slides were then scanned. Multiple ROIs were obtained per lesion, including necrotic core (when applicable), inner lesion, outer lesion border, and full thickness (encompassing inner and outer areas), as assessed by nuclear density and autofluorescence pattern. More fine-grained region determination was not possible due to poor performance of antibody staining. ROIs were then collected.

### GeoMx Library Prep and Sequencing

Following collection, GeoMx samples were removed from the machine and allowed to air dry overnight. The following day samples were placed in an open top thermal cycler at 65C for 10 minutes. Next, 10ul of nuclease free water were added to all samples well and pipetted up and down 5 times. PCR was run according to standard GeoMx protocols available in their quick start guide (MAN-10133-03). Pooling and cleanup were also run according to GeoMx protocols, with no deviations. The pooled library was assessed via Bioanalyzer and demonstrated a clean trace. Samples were loaded on the Illumina NextSeq platform at 1.6pM and sequenced twice using the recommended paired end 2 x 27 read acquisition. Sequenced library included 5% PhiX. Fastq files were assessed by QC metrics prior to further analysis.

### GeoMx data analysis

Raw probe counts from 2 sequencing runs were combined at the fastq levels and then converted to .DCC files via Nanostring’s geomxngspipeline function. The DCC files were uploaded to the DSP instrument and automatically associated with individual scans.

Sequencing and Probe QC was performed using default parameters (Analysis suite version 2.5.1.145). 4 of 32 original samples were removed for low sequencing saturation.

Because we intended to pseudobulk ROIs within the same animal and thus required normalization strategies not available on the DSP analysis suite, we exported two datasets: (1) raw, post-qc probe counts and (2) q3 normalized counts (the recommended normalization approach by Nanostring).

For normalization assessment, we first removed control probes from the non-normalized data. These raw data were assessed by PCA, which indicated strong biases induced by raw reads, surface area, and nuclei count, as expected. Given the publication of some biases that occur when using the Q3 normalization strategy on GeoMx data sets, we used a compositionally aware normalization strategy known as a centered-log ratio approach. To obtain CLR-transformed values for each gene, we first calculated the geometric mean of counts for each sample. We then created a ratio of an individual gene’s counts against the geometric mean from each sample. Finally, we calculated the log2 value of this ratio. Thus, all genes from a given sample were in essence normalized to their read-depth. Unlike proportional (relative) normalization strategies, this method preserves the opportunity for downstream statistical analyses. We next assessed the samples using their CLR-transformed data by PCA. The CLR normalization strategy effectively eliminated the relationships between PCA dimensions and read depth, surface area and nuclei count.

Interestingly, when compared to PCA based on the Q3 normalized data (which also directly accounted for surface area and nuclei), the results were highly concordant. This contrasts sharply with recent accounts of Q3-induced skew in GeoMx data sets, which our assessments indicate were the result of using small, focused gene panels like the Cancer Transcriptome Atlas – and not a fundamental flaw in the Q3 strategy. We have reached similar conclusions when using targeted gene sets on the Nanostring nCounter. Because this PCA analysis appeared to validate our use of the compositionally aware CLR approach, all downstream data used CLR values.

To create a pseudobulked data set, we first aggregated all raw, QC-counts originally exported from the DSP, which created 2 sets of data per animal: aggregated counts from granuloma-associated ROIs and counts from distal, uninvolved regions. Due to the removal of samples for low sequencing saturation (see above) 14/16 potential pseudobulk samples remained. Because CLR-transformed values are more appropriately assessed by Aitchison-distance PCoA (which is the Euclidean distance between CLR-transformed samples), PCoA was used for dimensionality reduction. To perform GSEA, we first calculated log2fc (using the CLR-transformed values) by directly comparing counts between necrotic and non-necrotic granulomas. We then used these log2fc values to rank genes. Ranked gene lists were supplied to a gsea function in R and results for significant enrichment and associated P values were obtained using the C5 Ontology gene sets from MSigDB.

### Luminex

For Luminex analyses, lungs from Mtb-infected mice were divided into 3 samples: the left lobe was homogenized in 1 ml PBS-Tween for CFU analysis, the inferior right lobe was placed in 5 ml Cytofix (BD) solution for overnight fixation and subsequent image analysis, and the remainder of the right lung was homogenized in 1 ml ProcartaPlex Cell Lysis Buffer (ThermoFisher) supplemented with Halt Protease Inhibitor (Invitrogen) and DNaseI (30 μg/ml; Sigma-Aldrich) to generate protein lysates. After homogenization, the lysate was pelleted at maximum speed at 4°C for 10 min, and the supernatant was centrifuged through two sequential rounds of 0.2 μm SpinX (Costar) columns to sterilize the sample for removal from the BSL3 facility. Homogenates were then assayed for protein levels using a custom 17-plex ProcartaPlex kit following the manufacturer’s instructions (Luminex). Homogenates were also assayed for total protein content using a BCA assay (Pierce), and protein levels of each analyte were normalized to 100 ug protein input.

### Single-cell RNA-sequencing

Single cell suspensions were generated from lung samples as described above prior to Mtb infection and at days 10, 17 and 34 post-Mtb infection. Cells were resuspended in 200 μl MACS buffer (PBS containing 2.5% FBS plus 1 mM EDTA), filtered through a 70 μm filter, and run on a FACS AriaII (BD) sorter. To collect parenchymal cells for single-cell RNA sequencing, alveolar macrophages (AM, SiglecF+CD11c+) were sorted separately into one collection tube to account for autofluorescence in the IV label channel, and all other IV-negative cells were sorted into another collection tube. After sorting, the two populations were combined and counted on a hemocytometer. After one round of washing with ice-cold DPBS, cells were resuspended to 1000 cells/μl in DPBS, and 8000 cells were inputted into the 10X Genomics pipeline following the manufacturer’s recommendations. After the generation of cDNA following the manufacturer’s protocol, samples were centrifuged through two sequential rounds of 0.2 μm SpinX (Costar) columns to sterilize the sample for removal from the BSL3 facility and subsequent library generation. Libraries were submitted to Psomagen (Rockville, MD) for NovaSeq sequencing, with 300M reads per sample.

### Alignment and processing of single cell RNAseq data

10X Chromium 3’ derived single-cell RNAseq sequence reads were aligned to the 10X Genomics pre-built mouse reference genome mm10-2020-A, assigned to individual cells by barcode, and UMI summarized using the 10X Cell Ranger 7.1.0 software package.

The Seurat R package was used for initial QC filtering and integration. First, a filtering step was applied across all samples, requiring all passing cells to have UMIs mapped to at least 500 distinct genes, and fewer than 5% of UMIs mapped to mitochondrial genes. Genes detected in fewer than 3 cells per mouse were excluded from further analysis. The Seurat integration pipeline^56^ was then applied to correct for batch effects and align cells across conditions including all combinations of mouse strain, Mtb strain, time post challenge, and CoMtb status.

Initial cell type assignment was performed using the CellTypist python package.^57^ As CellTypist does not have an available cell type model suitable for mouse lung or mouse immune cells, we created a de novo mouse lung immune cell type model using two published mouse cell atlases, namely the Tabula Muris^58^ and scMCA^59^ resources. Cell type labels were harmonized between both sources (e.g. macrophage -> Macrophage) and both datasets were filtered to retain immune and lung-associated cell types, excluding cells specific to other organs. The CellTypist python package was then used to train a mouse lung cell type model based on this combined resource. This model was then used to assign cell types to count-normalized log transformed data on a per-cell level from mouse lung scRNAseq samples, using the python scanpy^60^ package to normalize total counts per cell to 10,000 and log transform as required by CellTypist.

After initial cell type labelling, further unsupervised clustering of specific cell subtypes was performed, for cells labelled as ‘T cells’ or ‘NK cells’ and separately for all antigen presenting cell subtypes, i.e. “Alveolar macrophage”, “Dendritic cell”, “Monocyte” and “Macrophage”.

Unsupervised clustering was run using the standard Seurat pipeline which identifies the top 2,000 most variable genes in the data, creates a shared nearest-neighbor (SNN) network of cells, and divides the SNN into discrete clusters using the Louvain algorithm. The resulting clusters were manually annotated by identifying differentially expressed marker genes for each cluster (using the Seurat FindAllMarkers) function, and linking these marker genes to known cell types, e.g. Cd4+ IFNg+ Th1 cells express high levels of Cd3, Cd4 and Ifng).

To quantify changes in cell type proportion over time, total numbers of cells per-sample were calculated and normalized to cells per thousand per sample. Negative-binomial linear models, appropriate for zero-inflated count data, were fit and used to calculate p-values using the R glm.nb function.

Gene expression changes within specific cell types were determined using a pseudobulk approach, where counts from all similarly labelled cells were combined into a single sample x gene count matrix using the Seurat AggregateExpression function. The standard bulk RNAseq analysis package DESeq2^61^ was then used to calculate differential expression fold-changes and p-values for contrasts of interest.

Ranked gene lists from the above pseudobulk analysis were used as input for gene-set enrichment analysis using the R fgsea package.^62^ Gene sets used were previously-published human coherent blood transcriptional modules^63,64^ as available in the R tmod^65^ package, as well as mechanistic pathway modules from the REACTOME database^66^ as available in the R msigdbr package.^67^ To adapt human blood transcriptional gene sets to mouse, human genes were mapped to mouse orthologs using the Jackson Lab Mouse Genome Informatics Human-Mouse mapping [https://www.informatics.jax.org/downloads/reports/HOM_MouseHumanSequence.rpt]. The resulting mouse-translated gene sets were filtered to retain only blood transcriptional modules with at least 5 mouse genes where > 80% of the original human genes were successfully mapped to mouse orthologs. Unannotated gene sets (“TBA” or “Undetermined”) were removed from further analysis.

We used the CellChat analysis package^28^ (version 1.6.1) to quantify the strength of receptor-ligand communications among cell types in our scRNAseq dataset. To simplify interpretability and ensure a sufficient number of cells of each type in the analysis, the sub-types of CD4+ and CD8+ T cells were grouped into the broader categories “CD4+ T cell” and “CD8+ T cell”. Additionally, the IM and monocyte sub-types were grouped into “monocyte-derived cells” and AM sub-types were grouped into a broader “AM” category. Our analyses focused on examining 1) whether intercellular communications originating with T cells and targeting myeloid cells differed between the primary and CoMtb conditions and 2) whether neutrophil-to-neutrophil chemotactic communications (those represented in the “CCL” and “CXCL” CellChat pathways) differed between conditions.

### Quantification and statistical analysis

Statistical tests were selected based on appropriate assumptions with respect to data distribution and variance characteristics. Statistical details of experiments can be found in the figure legends. No statistical methods were used to predetermine sample size. The statistical significance of differences in mean values was determined by the appropriate test, as denoted in the figure legends. Paired t tests were performed only when comparing responses within the same experimental animal or tissue, or group means within the same experiment (indicated in the legend). Correlations and corresponding p values by Pearson’s correlation test. ∗∗∗∗, p ≤ 0.0001; ∗∗∗, p ≤ 0.001; ∗∗,p ≤ 0.01; and ∗, p ≤ 0.05; NS, p > 0.05.

## Supplemental Information

**Figure S1. Related to Figure 1. Pre-existing immunity abrogates the formation of necrotic granulomas:** A) Lesion zoom-ins from Fig 1A. B) PCA loadings from Fig 1B. C) Representative confocal images showing necrotic debris in center of primary necrotic lesion, and intact nuclei of immune cells within lesion formed in setting of CoMtb. D) Histocytometry gating scheme to determine cell types for analysis in Figs 1G-K.

**Figure S2. Related to Figure 2. CoMtb alters the immune landscape following Mtb infection, and Figure 3: CoMtb accelerates T cell and MDC activation, blunts neutrophil responses:** A) Top 50 DEGs for necrotic and non-necrotic lesions in Fig 2A. B) Top genes which discriminate scRNAseq clustering into the specific cell types in Fig 2C. C) Correlations of cell proportion as determined by scRNAseq with CFU. D) Early and late neutrophil clustering and top 10 DEGs, corresponding to Figs 2F,G. E) GSEA analysis comparing CoMtb to primary Mtb infection across timepoints, corresponding to Fig 3A. FDR determined by the R fgsea package. Correlations by Pearson’s correlation test.

**Figure S3. Related to Figure 2. CoMtb alters the immune landscape following Mtb infection, and Figure 3: CoMtb accelerates T cell and MDC activation, blunts neutrophil responses:** A) Pulmonary bacterial burdens corresponding to Fig 2E. B) Lymphoid flow cytometry gating scheme for Fig 2E. C) Myeloid flow cytometry gating scheme for Fig 2E. D) Pulmonary bacterial burdens corresponding to Fig 3D.

**Figure S4. Related to Figure 5. CD4 T cells are required for CoMtb-mediated protection from lesion necrosis:** A) Representative flow plots and CD4 T cell enumeration following □CD4 depleting antibody administration. B) Representative flow plots and neutrophil enumeration following □CD4 depleting antibody administration. Single-group comparisons by unpaired t test.

**Figure S5. Related to Figure 6: Neutrophils drive lesion necrosis:** A) Representative flow plots and neutrophil enumeration following □Ly6G depleting antibody administration. Also demonstrates concordance of CD177 and Ly6G in an Mtb-infected lung. B) Heatmap showing cellular composition of clustered microenvironments and percent area of lesion comprised by each microenvironment, uninvolved regions not included, corresponding to Fig 6I. C) Heatmap showing cellular composition of clustered microenvironments and percent area of lesion comprised by each microenvironment, uninvolved regions not included, corresponding to Fig 6N. D) Confocal microscopy image of one small necrotic lesion with low antigen abundance, identified following Late Ly6G depletion.

**Table S1: Related to Figure 1. Pre-existing immunity abrogates the formation of necrotic granulomas:** Pathology scores for hematoxylin and eosin-stained tissue sections in Figure 1A.

