## Supplementary figures and images for "CD4-mediated immunity shapes neutrophil-driven tuberculous pathology"

### Supplemental Figure 1

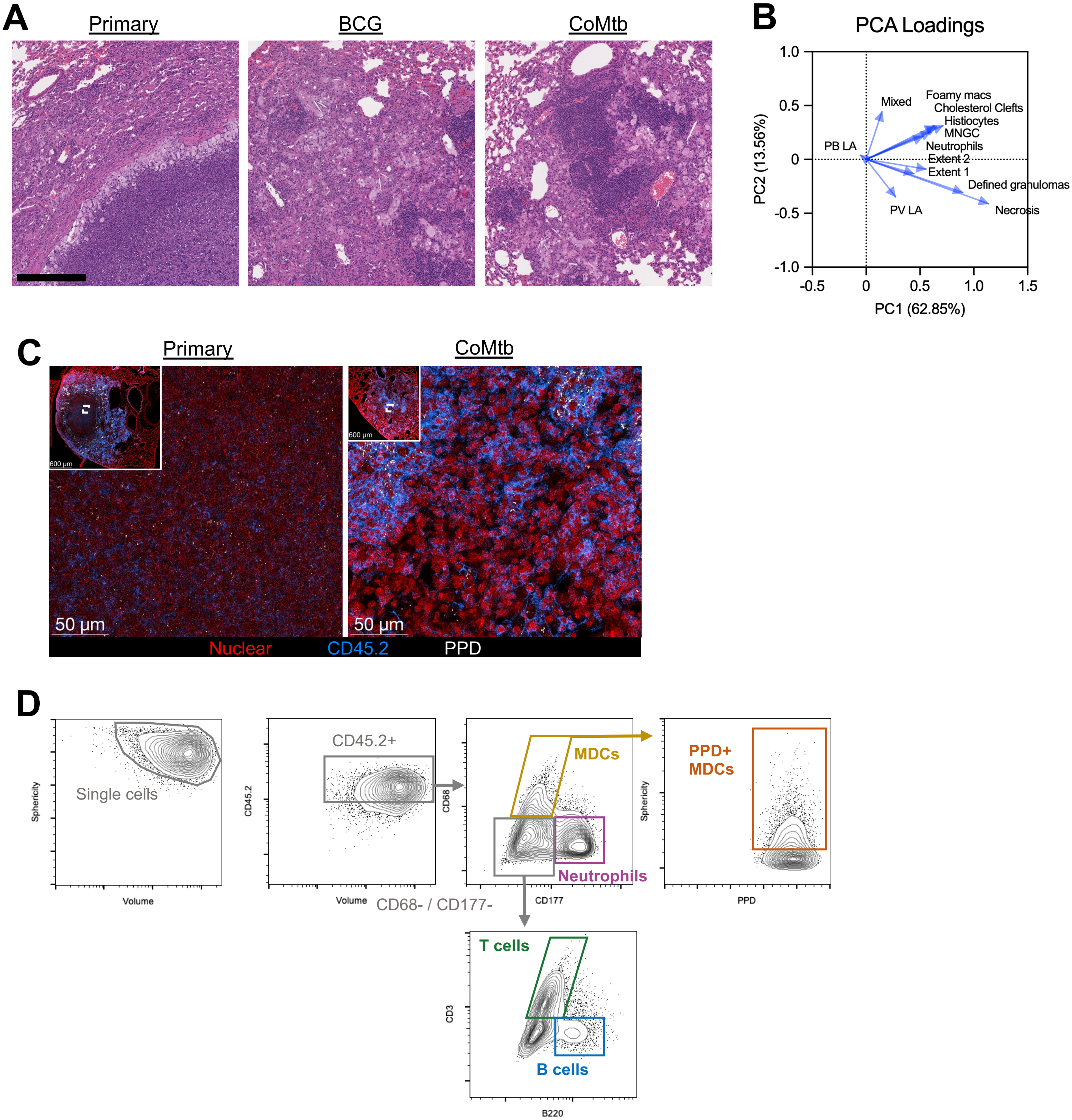

### Supplemental Figure 2

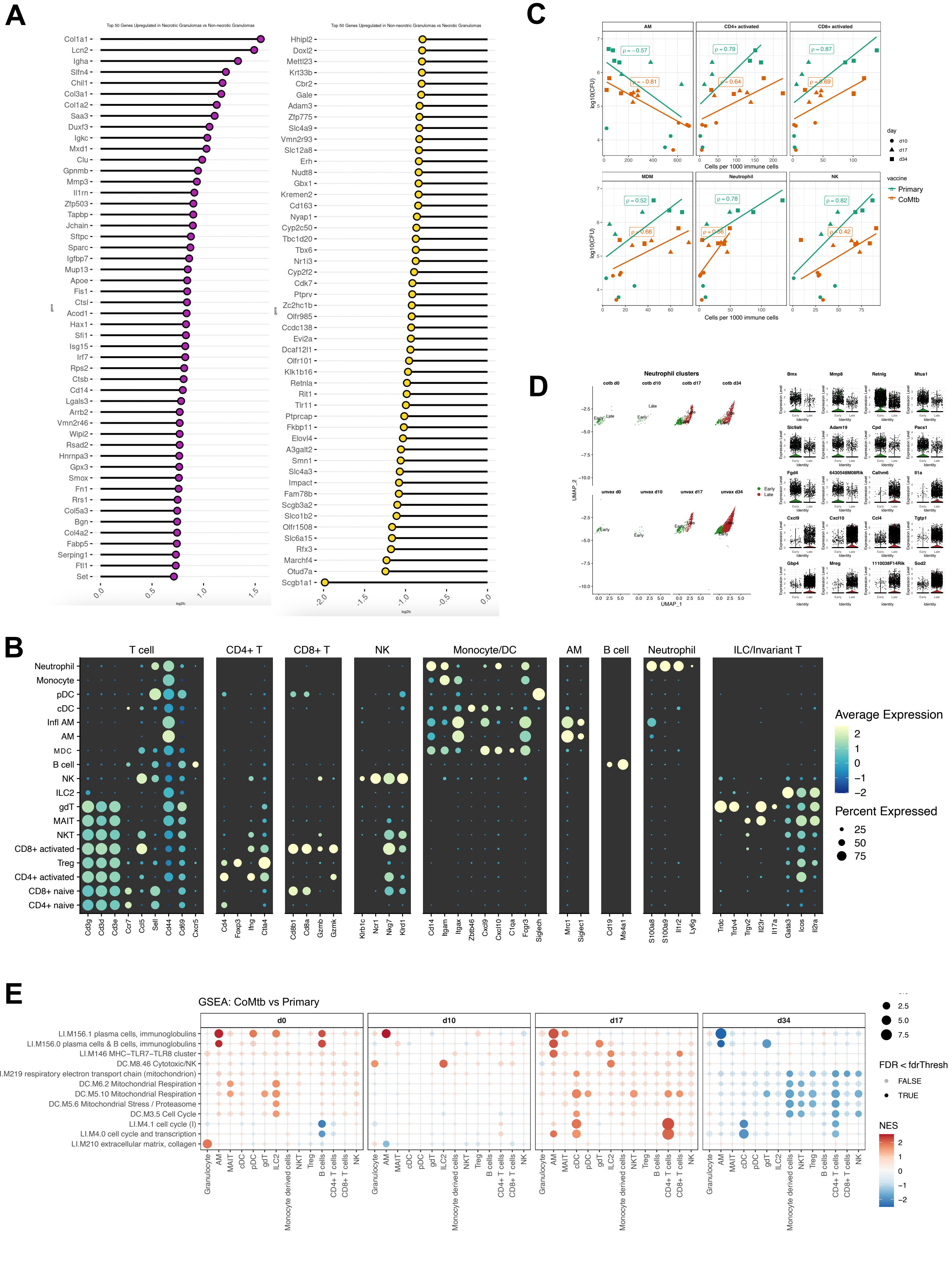

### Supplemental Figure 3

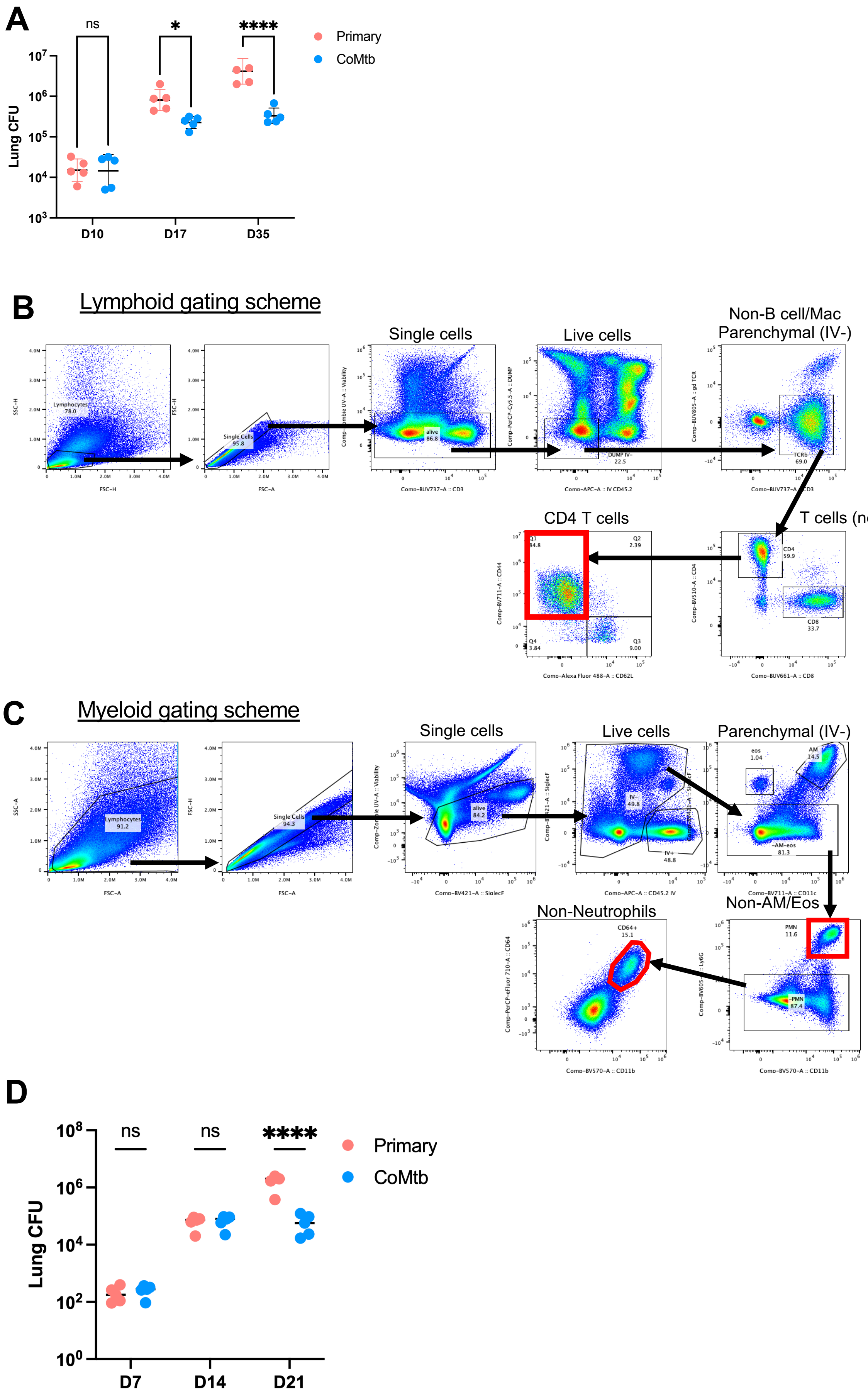

### Supplemental Figure 4

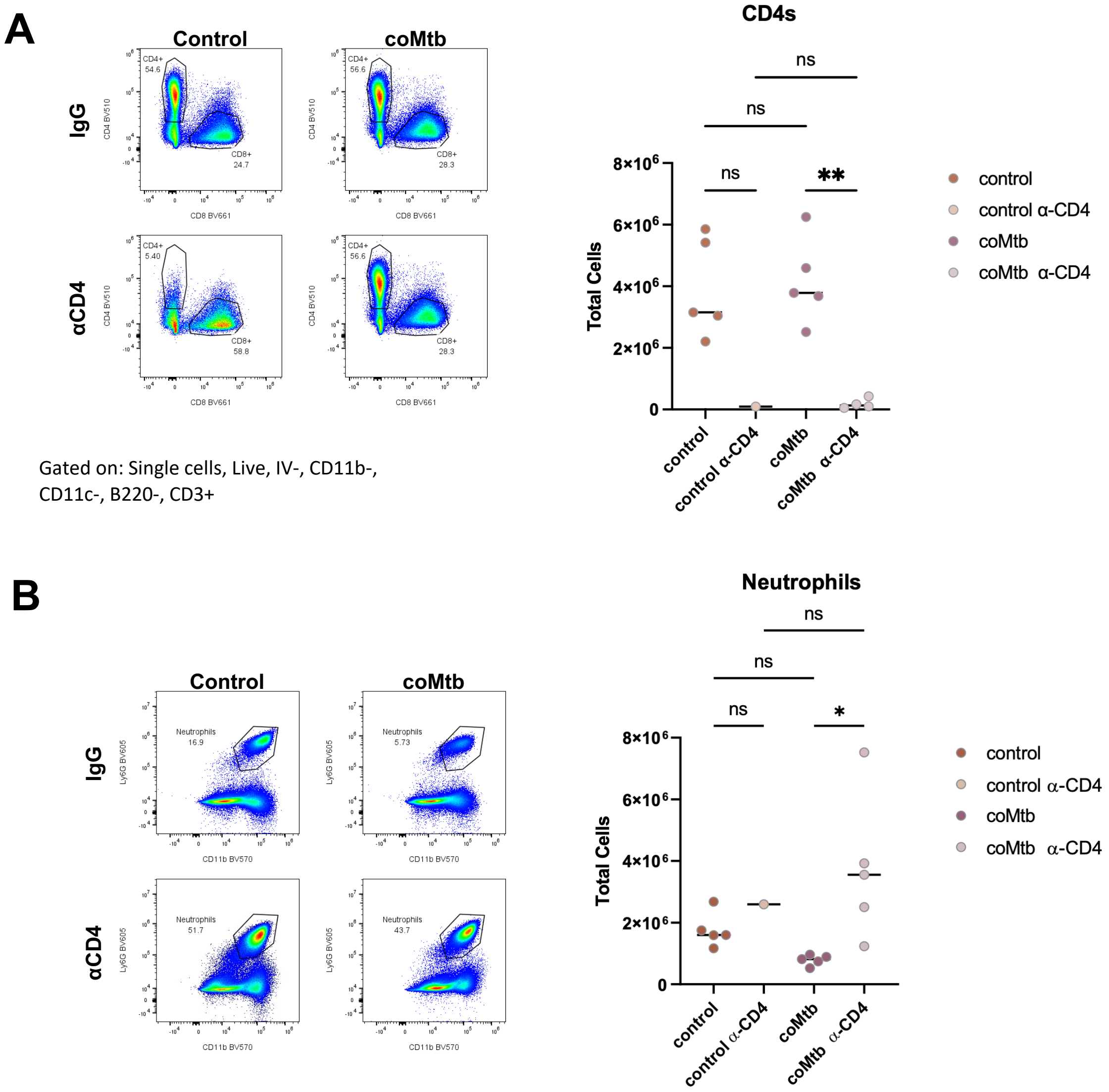

### Supplemental Figure 5

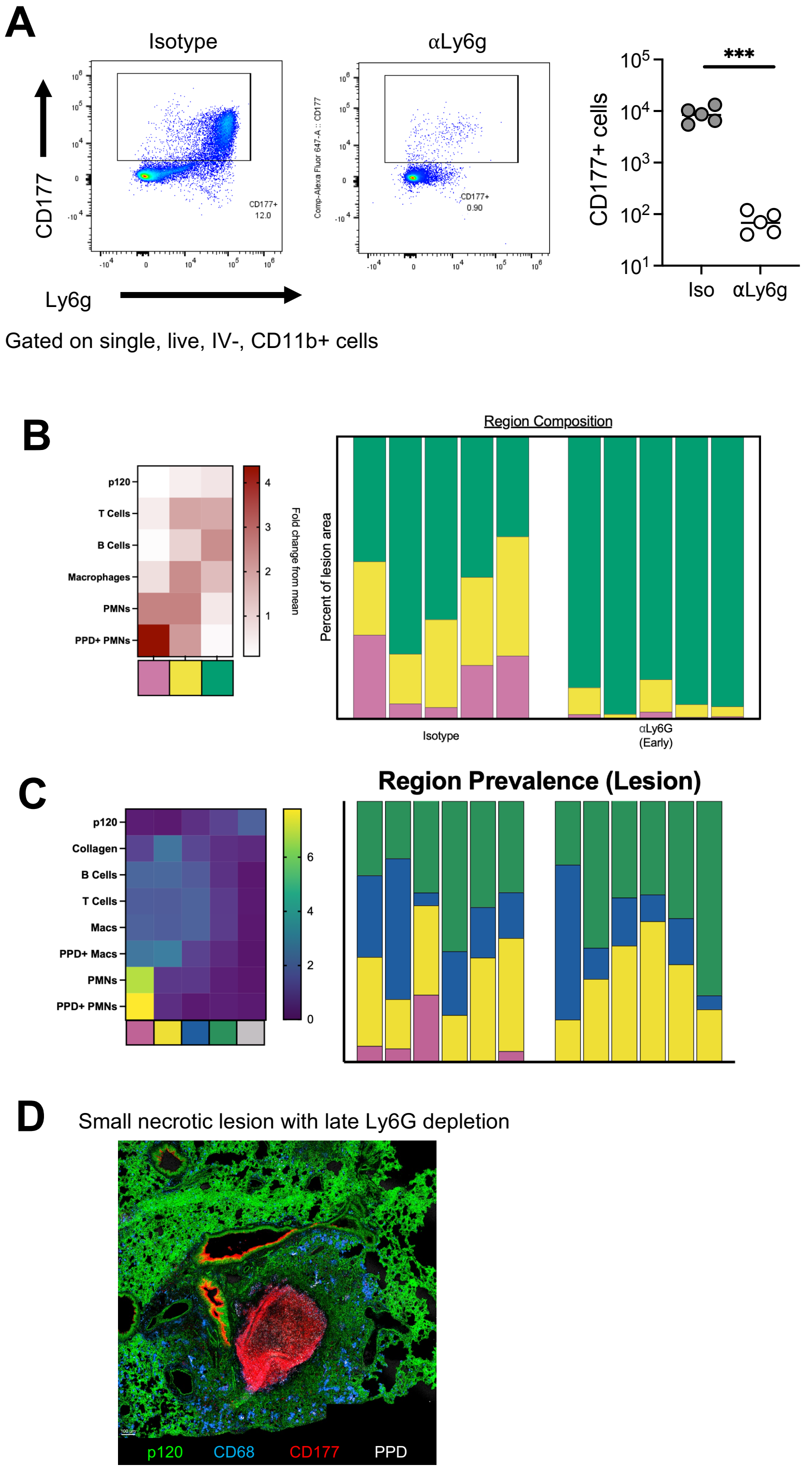
