## Supplemental Table 1 for "CD4-mediated immunity shapes neutrophil-driven tuberculous pathology"

|  | **Extent 1** | **Extent 2** | **Mixed Granulomas** | **Defined Granulomas** | **Perivasclar Lymphoid Aggregates** | **Peribroncial Lymphoid Aggregates** | **Histiocytes** | **Foamy Macrophages** | **Multinucleated Giant Cell** | **Neutrophils** | **Necrosis** | **Cholesterol Clefts** |
| --- | --- | --- | --- | --- | --- | --- | --- | --- | --- | --- | --- | --- |
| **Primary 1** | 3 | 3 | 1 | 4 | 2 | 1 | 3 | 3 | 2 | 2 | 3 | 2 |
| **Primary 2** | 3 | 2 | 2 | 2 | 2 | 1 | 3 | 3 | 2 | 3 | 1 | 2 |
| **Primary 3** | 3 | 3 | 1 | 3 | 2 | 1 | 3 | 3 | 1 | 2 | 3 | 2 |
| **Primary 4** | 3 | 2 | 1 | 3 | 2 | 0 | 3 | 3 | 1 | 2 | 3 | 2 |
| **Primary 5** | 4 | 3 | 1 | 4 | 3 | 1 | 3 | 3 | 2 | 2 | 4 | 2 |
| **BCG 1** | 3 | 2 | 1 | 2 | 2 | 1 | 3 | 4 | 1 | 2 | 1 | 2 |
| **BCG 2** | 2 | 2 | 2 | 2 | 2 | 1 | 3 | 3 | 1 | 2 | 1 | 2 |
| **BCG 3** | 3 | 2 | 2 | 2 | 1 | 1 | 3 | 3 | 2 | 2 | 1 | 2 |
| **BCG 4** | 2 | 2 | 1 | 2 | 1 | 1 | 2 | 2 | 1 | 1 | 1 | 2 |
| **BCG 5** | 3 | 2 | 1 | 3 | 1 | 1 | 3 | 3 | 1 | 2 | 1 | 2 |
| **CoMtb 1** | 2 | 1 | 0 | 1 | 1 | 1 | 1 | 1 | 0 | 1 | 0 | 0 |
| **CoMtb 2** | 2 | 1 | 2 | 1 | 1 | 1 | 2 | 2 | 1 | 0 | 1 | 1 |
| **CoMtb 3** | 3 | 1 | 1 | 1 | 3 | 1 | 1 | 1 | 0 | 1 | 0 | 0 |
| **CoMtb 4** | 2 | 2 | 0 | 3 | 2 | 1 | 2 | 3 | 0 | 1 | 0 | 1 |
| **CoMtb 5** | 2 | 2 | 1 | 1 | 1 | 1 | 2 | 2 | 0 | 1 | 0 | 0 |
